# Network propagation of rare mutations in Alzheimer’s disease reveals tissue-specific hub genes and communities

**DOI:** 10.1101/781203

**Authors:** Marzia A. Scelsi, Valerio Napolioni, Michael D. Greicius, Andre Altmann, for the Alzheimer’s Disease Neuroimaging Initiative (ADNI), the Alzheimer’s Disease Sequencing Project (ADSP)

## Abstract

**Background:** State-of-the-art rare variant association testing methods aggregate the contribution of rare variants in biologically relevant genomic regions to boost statistical power. However, testing single genes separately does not consider the complex interaction landscape of genes, nor the downstream effects of non-synonymous variants on protein structure and function. Here we present the NETwork Propagation-based Assessment of Genetic Events (NETPAGE), an integrative approach aimed at investigating the biological pathways through which rare variation results in complex disease phenotypes.

**Results:** We applied NETPAGE to sporadic, late-onset Alzheimer’s disease (AD), using whole-genome sequencing from the AD Neuroimaging Initiative (ADNI) cohort, as well as whole-exome sequencing from the AD Sequencing Project (ADSP). NETPAGE is based on network propagation, a framework that models information flow on a graph and simulates the percolation of genetic variation through gene networks. The result of network propagation is a set of smoothed gene scores used to predict disease status through sparse regression. The application of NETPAGE to AD enabled the identification of a set of connected genes whose smoothed mutation profile acted as a robust predictor of case-control status, based on gene interactions in the hippocampus. Additionally, smoothed scores significantly correlated with risk of conversion to AD in Mild Cognitive Impairment (MCI) subjects. Lastly, we showed tissue-specific transcriptional dysregulation of the core genes in two independent RNA-seq datasets, as well as significant enrichments in terms and gene sets with known connections to AD.

**Conclusions:** The presented framework enables enhanced genetic association testing for a wide range of traits, diseases, and sample sizes.

## BACKGROUND

The advent of next generation sequencing (NGS) has drastically changed the genetic landscape of both complex and Mendelian traits, widening the range of approaches to investigate the genetic bases of human phenotypes. Using SNP-array genotyping technologies, large scale studies involving 100,000s of participants are focusing on common variants in order to identify loci associated with complex traits [1]. However, owing to natural selection, common variants typically do not impart major risk for disease [2], with rare exceptions of variants such as the ε4 allele of *APOE* in Alzheimer’s disease (AD) [3,4]. Loci identified in genome wide association studies (GWAS) are often flagging the existence of haplotypes harbouring rare variants with strong disease effects and therefore have to be followed up by fine-mapping to identify the true causal variants.

NGS, by contrast, is most effectively used when focusing on rare genetic variants, private mutations or structural genome changes. These types of rare variants have the possibility to exercise a large effect on disease risk and often show a Mendelian inheritance pattern, thus being at the core of familial forms of many disorders; prominent examples are mutations in *APP*, *PSEN1*, and *PSEN2* in familial AD [5] or variants in *MAPT*, *GRN* and *C9orf72* in frontotemporal dementia (FTD) [6]. The decreasing costs for NGS have enabled the establishment of large whole genome (WGS) or whole exome (WES) sequencing studies such as the AD Sequencing Project (ADSP) [7]; still the largest studies are two orders of magnitude smaller than the largest GWAS (i.e., 10,000s vs a million participants). Moreover, by definition, rare variants are not frequent in the population and it is unlikely for the same rare variant to be shared by many subjects in a study with limited sample size. Therefore, the resulting data matrix of subjects × rare variants is sparse. This drawback is often addressed with alternative study designs such as “extreme phenotyping”, based on the assumption that rare mutations accumulate in the extreme tails of the phenotype distribution.

Low sample size and sparsity pose a problem for analysing these types of genetic data: two determinants of the statistical power for classical association studies are sample size and allele frequency, leading to low statistical power in rare variant association analyses when relying on the typical single-variant GWAS methodology. This problem has motivated the development of advanced statistical methods, reviewed in [8]. The most straightforward approach is the gene-wise burden test, i.e., for each gene the number of rare nonsynonymous variants is counted per participant and the gene burden is compared between cases and controls. The underlying assumption, in order to convert a sparse matrix into a non-sparse one, is that affected individuals tend to carry mutated versions of key genes, while healthy individuals do not. This assumption has been proven to be correct in the case of HDL cholesterol levels [9]. This basic model has been superseded by methods based on the Sequence Kernel Association Test (SKAT) [10], which performs a variance-component test: this assumes that the genetic effect for a given variant (regression coefficient for the SNP in a mixed model) follows a distribution with mean 0 and variance 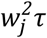, and then tests the null hypothesis *H_0_*: *τ* = *0*. One key advantage of this test is that the rare variants within the gene do not have to influence the disease risk in the same direction. SKAT has been successfully used to study rare variants in a number of diseases such as AD [11], schizophrenia [12], and health and disease more generally [13].

Still, SKAT and its extensions focus on predefined genetic regions, such as genes or even pathways. Recent observations, however, suggest that the underlying nature of genetic effects is more complicated. For instance, the recently proposed omnigenic model for complex traits [14] postulates that genetic variation percolates through gene interaction networks and that, due to the small-world property of such networks, any gene is only a few steps away from the “core” genes with specific roles in disease etiology, such that the effect of variation flows from peripheral genes to “core” genes. In fact, an exonic, deleterious variant within a gene by definition will lead to changes in the resulting protein (e.g., amino acid substitution, premature truncation of transcription, loss of a stop codon). These changes will in turn affect the protein’s downstream interactions within the cell environment. Ultimately this may disrupt one or more molecular pathways through a chain reaction (or domino effect). This small-world property implies that mutations in different, unrelated genes and different patients can “converge” and exert their effect on the same core gene (Figure 1). Furthermore, complex traits are mediated through multiple tissues or cell types and, consequently, the quantitative effect of genetic variation will vary across tissues owing to differences in gene and protein interaction networks.

**Figure 1.**
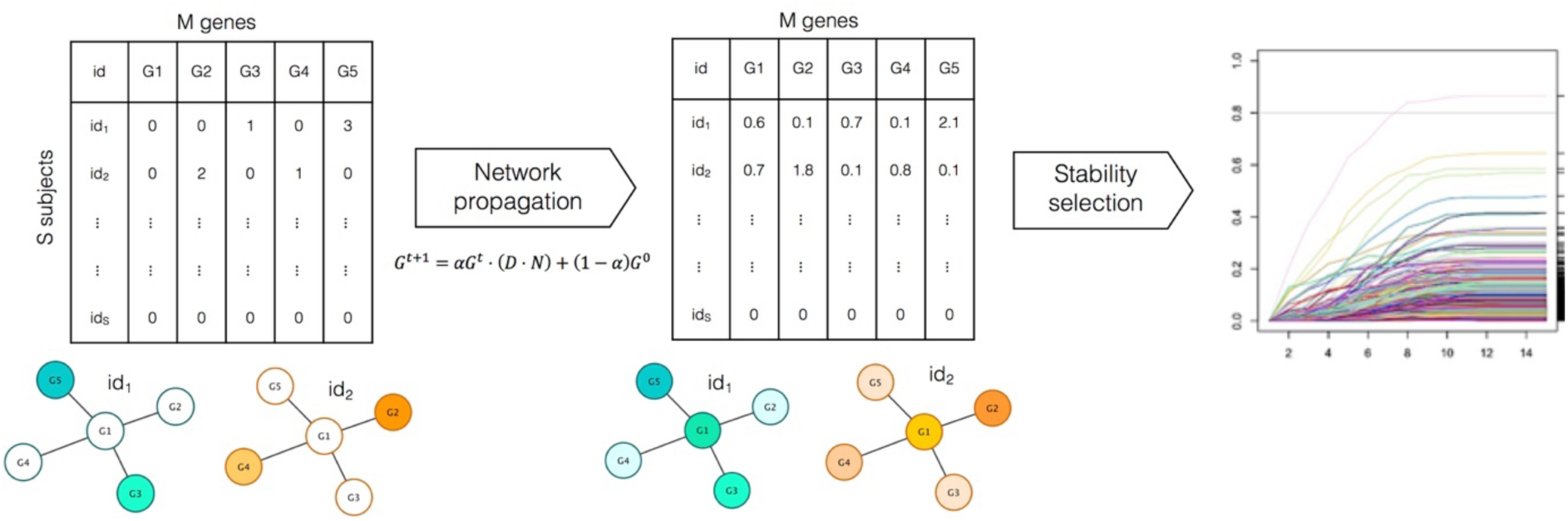
NETPAGE combines network propagation with sparse regression to follow the pathways by which rare mutations percolate through gene interaction networks. In the network propagation process, the flow of information through the network is controlled by the diffusion length α. The matrix G in the formula represents the mutational burden at iterations *t* and *t+1*; while D·N is a degree-normalised adjacency matrix of the gene interaction network (see Methods). The result of network propagation is a set of smoothed gene scores used to predict disease status through sparse regression and stability selection.

In this work we propose a gene-based test for rare variation whose rationale resembles the omnigenic assumption. In particular, we leverage the network propagation approach, which models a diffusion process or information flow on a graph structure, in order to delineate the percolation effect of genetic variation through gene networks. Additionally, we embed tissue specificity in our framework by using tissue-specific gene-interaction networks obtained from Greene *et al.* [15]. Network propagation is an established approach that has enabled methodological advances as well as important findings in many research fields. In particular, it has been defined as an “universal amplifier of genetic associations” [16]. In brief, network propagation has been successfully applied to various bioinformatics tasks, such as the prioritization of disease-associated genes based on gene interactions and disease similarities [17] or based on a modified version of Google’s PageRank algorithm [18], and the stratification of tumour subtypes based on somatic mutation signatures [19]. A more comprehensive review of applications of network propagation can be found in [16]. Notably, an early attempt at gene prioritisation based on tissue-specific interaction networks dates back to 2012 [20]. However, the networks developed for this work leveraged the tissue specificity of gene expression only, whereas Greene *et al.* [15] integrated a much wider variety of genomic data types, comprising gene co-expression, transcription factor regulation, protein interaction, chemical and genetic perturbations, and microRNA target profiles.

Here we present the NETwork Propagation-based Assessment of Genetic Events (NETPAGE), an integrative approach that combines network propagation with sparse regression in order to investigate the biological pathways through which genetic variation affects tissue function and results in complex disease phenotypes. Our approach is highly generalisable and enables enhanced genetic association testing for a wide range of complex traits (binary and continuous), diseases, and sample sizes. As a specific proof of concept, we applied NETPAGE to study rare genetic variation in a small whole genome sequencing (WGS) dataset focusing on sporadic late-onset AD obtained from the AD Neuroimaging Initiative (ADNI), as well as a medium-sized whole exome sequencing (WES) dataset from the AD Sequencing Project (ADSP).

## RESULTS

### Method overview

NETPAGE combines network propagation with sparse regression to identify genes robustly associated with a phenotype (Figure 1). In this work we focused on rare, exonic, deleterious Single Nucleotide Variants (SNVs) from WGS or WES; different criteria for the selection of rare deleterious SNV were utilised, to assess their impact on the final result (see Methods for details). SNVs were projected onto a gene interaction network. We compared the hippocampus network derived by Greene et al. [15] with the whole, non-tissue-specific human interactome available in STRING [21]. Network propagation is then used to model the propagation of the effects of rare mutations through the network. Network propagation results in a set of gene-wise, continuous “smoothed” scores that are then tested for robust association with a disease phenotype through sparse regression (LASSO [22]) and stability selection [23]. NETPAGE is not intended as a classification tool, hence we did not investigate its prediction performances in the classic machine-learning sense.

### Network propagation: simulations

In order to explore the effect of different parameters in the network propagation algorithm we conducted a series of simulations to map out reasonable default parameters for the algorithm. The key parameters of network propagation are the distance a mutation signal is allowed to travel through the network (termed diffusion length α from now on), and the percentage of top edges to be retained in the network in a procedure called binarisation. We generated synthetic datasets, varying parameters of mutation frequency in cases and controls, and applied network propagation using varying parameter settings of the algorithm. In these simulations we focus on an un-mutated target gene and vary the mutation frequency in the gene’s neighborhood differentially between cases and controls. Next, we test how well the smoothed score obtained through network propagation in the target gene separates cases from controls. Further details can be found in SI Appendix. A summary of results from the investigation of the parameter space for network propagation on simulated data can be found in Figure 2 and Supplementary Figure 1. We observed (Supplementary Figure 1B) that variation in both diffusion length α and percentage of edges retained influences the difference in a hub gene’s smoothed score between cases and controls only marginally. Therefore we adopted a parsimonious approach and selected as default parameters for real data applications a mid-range α value of 0.5 and a top edge percentage of 1%. As expected, for α=0 no propagation occurs. We also observed (Supplementary Figure 1A) a detrimental effect on the smoothed scores (loss of statistical significance) when quantile-normalisation, which is used in Hofree et al. [19], is applied at the end of network propagation, therefore we decided to exclude this step from our pipeline.

**Figure 2.**
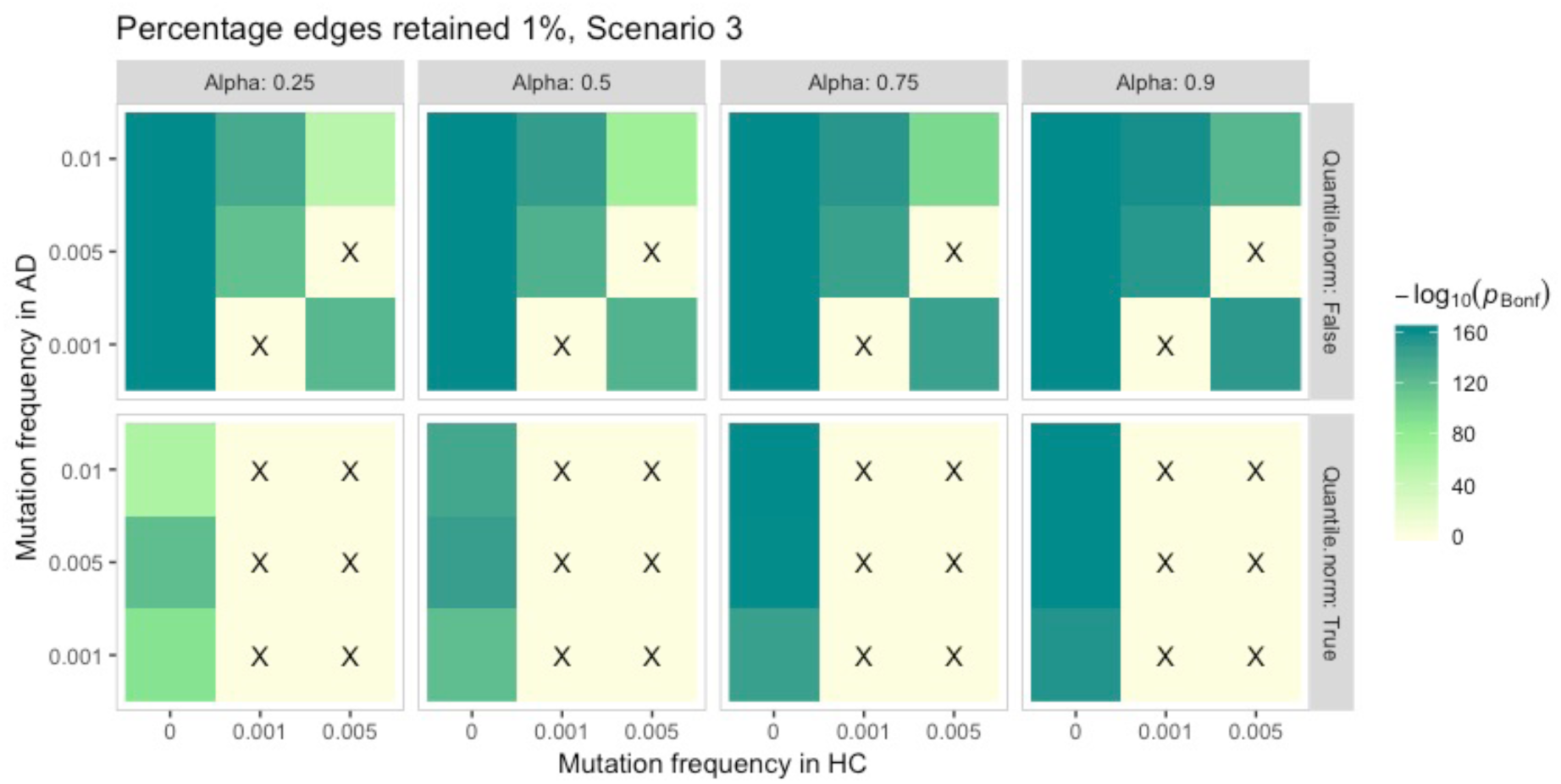
A selection of simulation results. The x and y axis represent the mutation frequencies in controls and cases respectively. The faceting allows to visualise the effect of other parameters (diffusion length, quantile normalisation). The colour-coding indicates the statistical significance of the difference in the hub gene smoothed score between controls and cases, in units of −log10 p-value (Bonferroni corrected). We investigated three mutation scenarios: scenario 1, only first neighbours mutated; scenario 2, only second neighbours mutated; scenario 3, both first and second neighbours mutated; the target gene is always unmutated. Cells marked with a black cross indicate parameter combinations where the smooth score of the target gene was not significantly different between cases and controls. This set of tile plots shows the effect of varying diffusion length quantile normalisation after network propagation with top 1% edges retained in scenario 3.

### NETPAGE: application to AD sequencing data

After verifying that the network propagation step alone works as expected, we applied the complete NETPAGE framework to two independent, real-world NGS datasets: one with a small sample size (ADNI), and another with a medium sample (ADSP).

#### Small sample: ADNI

We used NETPAGE in ADNI to test smoothed scores for M = 13,310 genes for association with binary disease outcome in N = 439 Caucasian subjects (222 healthy controls [HC], 217 AD). One gene resulted from stability selection on the smoothed scores in ADNI (Figure 3A): *PFAS* (selection probability = 0.85; Table 2). We replicated the selection of PFAS when testing the gene burden propagated through the STRING network (selection probability = 0.85; Supplementary Figure 2). No genes were selected when running stability selection on the “raw” mutation profile (α = 0, no smoothing), nor when the mutation profile was smoothed through either a randomised version of the hippocampus network or a non-brain-related network (umbilical cord; Supplementary Figure 3).

**Figure 3.**
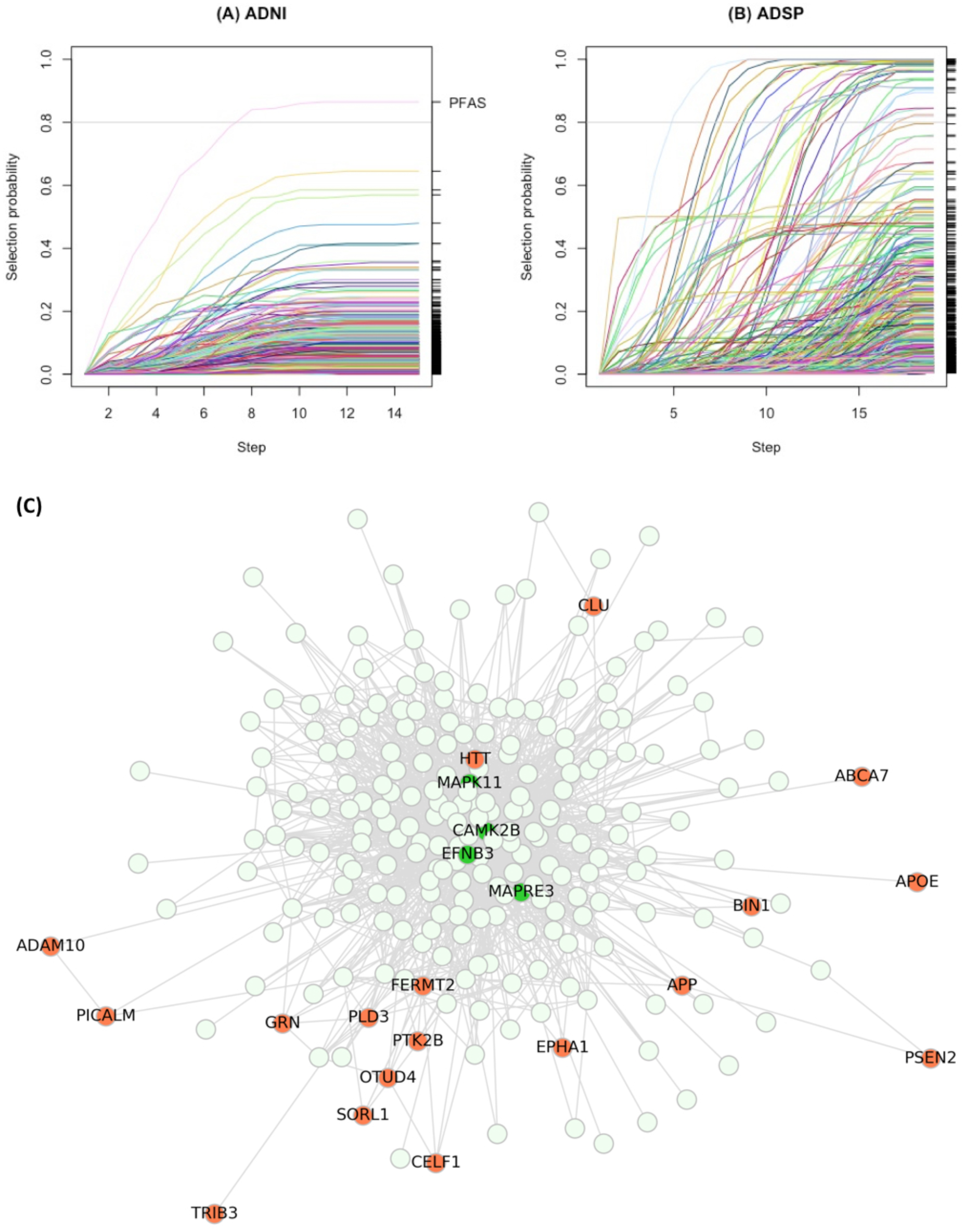
Stability selection results in ADNI (A) and ADSP (B), for a selection probability cutoff of 0.80. In these stability paths plots, the x axis represents the steps taken along the lambda sequence to choose the optimal amount of regularisation required by the LASSO; the y axis represents the selection probability of a gene. Selection probability paths (trajectories) for different genes are represented by different colours. Genes whose trajectories crossed the threshold of 0.80 selection probability were considered as robust predictors of case-control status and followed up in subsequent analyses. Panel (C) shows the ego network of *CAMK2B* (selected in ADSP). Nodes coloured in bright green were identified by NETPAGE; nodes coloured in coral red are genes linked to AD and other neurodegenerative disease in the literature.

When a burden including only rare deleterious stop-gain, stop-loss and frameshift mutations was smoothed (through either the hippocampus or the STRING PPI network) and tested for association with diagnosis, no genes were selected (Supplementary Figure 4).

#### Medium sample: ADSP

We used NETPAGE in ADSP to test smoothed scores for M = 16,298 genes for association with binary disease outcome in N = 10,186 Caucasian subjects (4601 HC, 5585 AD). Stability selection on the smoothed scores in ADSP identified 29 genes as robust predictors of case-control status (Figure 3B). Selection probabilities for these 29 genes are reported in Table 2. As an example, figure 3C shows the subgraph of radius two centred on *CAMK2B*, including interesting second neighbours of relevance to AD dementia or other neurodegenerative disorders. Eighteen genes selected from ADSP were found to be second neighbours of the gene selected from ADNI (*PFAS*) in the hippocampus network. This number is significantly higher than it would be expected by chance (p < 10^-4^; Supplementary Figure 5), indicating that NETPAGE-selected genes in ADSP are overrepresented in the interactome of PFAS.

As common variants identified through GWAS are often tagging loci harbouring rare variants, Supplementary Table 1 lists selection probabilities for the 21 genes reported in a recent large scale GWAS for AD [24].

### Gene-based rare variant association testing

NETPAGE was compared to SKAT-O [25], a state-of-the-art method for gene- and set-based association testing of rare variants. Associations with diagnosis were tested for N = 439 Caucasian ADNI subjects and M = 13,591 genes controlling for age, sex, number of *APOE* ε4 alleles, years of education, and population structure (see Methods). Additionally, to conduct a fair comparison to SKAT-O, a mass univariate test of association between the smoothed gene scores and case-control status was also carried out via logistic regression for M = 13,310 genes, controlling for the same confounders listed above. After correction for multiple comparisons, no genes were significantly associated with case-control status in either test (SKAT-O, Supplementary Figure 4A; mass univariate smoothed, Supplementary Figure 4B).

We also sought to compare SKAT-O and NETPAGE on a dataset where SKAT-O would be sufficiently powered to detect associations [11]. We therefore grouped 270,165 non-synonymous SNVs from 10,186 Caucasian ADSP samples into 16,630 genes and performed a gene-based test, correcting for age, sex and number of APOE4 alleles. After correction for multiple comparisons, no genes were significantly associated with case-control status in ADSP (Supplementary Figure 6).

### Set-based rare variant association testing

Recent research has shown that the cumulative effect of rare deleterious variation in relation to disease can in some cases be revealed by simply aggregating SNVs into genes and then pathways when running SKAT [26]. To demonstrate the added value of the network propagation step over this approach, we conducted set-based association testing for rare, deleterious SNVs with case/control status in ADNI. Gene sets were defined from the results of stability selection: each selected gene was grouped with its first neighbours in the hippocampus network. Supplementary Table 2 reports association p-values with case-control status in ADNI for (A) burden, (B) variance-component, and (C) omnibus test (SKAT-O) for rare variants in the gene sets defined around selected genes. The gene grouping around *PFAS* was not associated with case-control status.

### Model comparison

The selection of genes whose smoothed scores correlated with case-control status was performed without considering the relative contribution of other established confounders such as sex and age. Therefore we sought to assess whether or not the smoothed scores carry any additional predictive power on top of the variation captured by such confounders, and to what extent. In ADNI, an extended logistic regression model including the smoothed score for *PFAS* significantly improved goodness-of-fit to case-control status over a baseline model including established AD predictors (sex, age, APOE ε4 count, education, population structure; chi-squared test p = 2×10^-4^). Additionally, the extended model showed a higher pseudo-R^2^ statistic (p-R^2^ = 0.36) than the baseline model (p-R^2^ = 0.33), approximately equivalent to an additional 3% variance explained in the outcome.

In ADSP, we found that *CSNK1A1* was a second neighbour of *PFAS*, had a selection probability of 1 and provided the best improvement in goodness-of-fit to case-control status over the baseline model including sex, age and APOE ε4 (chi-squared test corrected p = 1×10^-36^; Table 2).

### Survival analysis

We postulated that the smoothed scores possess unique properties in that they condense in a biologically meaningful way information on how mutations in a neighbourhood interact, acquiring an increased sensitivity to disease phenotypes. We therefore set out to further characterise the properties of such scores by assessing their relationship not only to simple case-control status but also to risk of clinical disease progression to AD.

The **“**raw” (binary) mutation status and smoothed score for *PFAS* in ADNI were tested for association with risk of conversion to AD using Cox proportional hazards model. The binary mutation status of *PFAS* was not seen to influence the probability of conversion to AD (p = 0.65; Supplementary Figure 7A). The inclusion of covariates (see Methods) confirmed this result (pPFAS = 0.82; Supplementary Figure 7B). Conversely, the smoothed score of *PFAS* showed a statistically significant protective effect against conversion to AD (pPFAS < 0.001; Figure 4) on top of the effect of other established confounders. The score was still significantly associated when removing healthy controls from the survival analysis to partially avoid circularity issues (p = 0.008; Supplementary Figure 7C).

**Figure 4.**
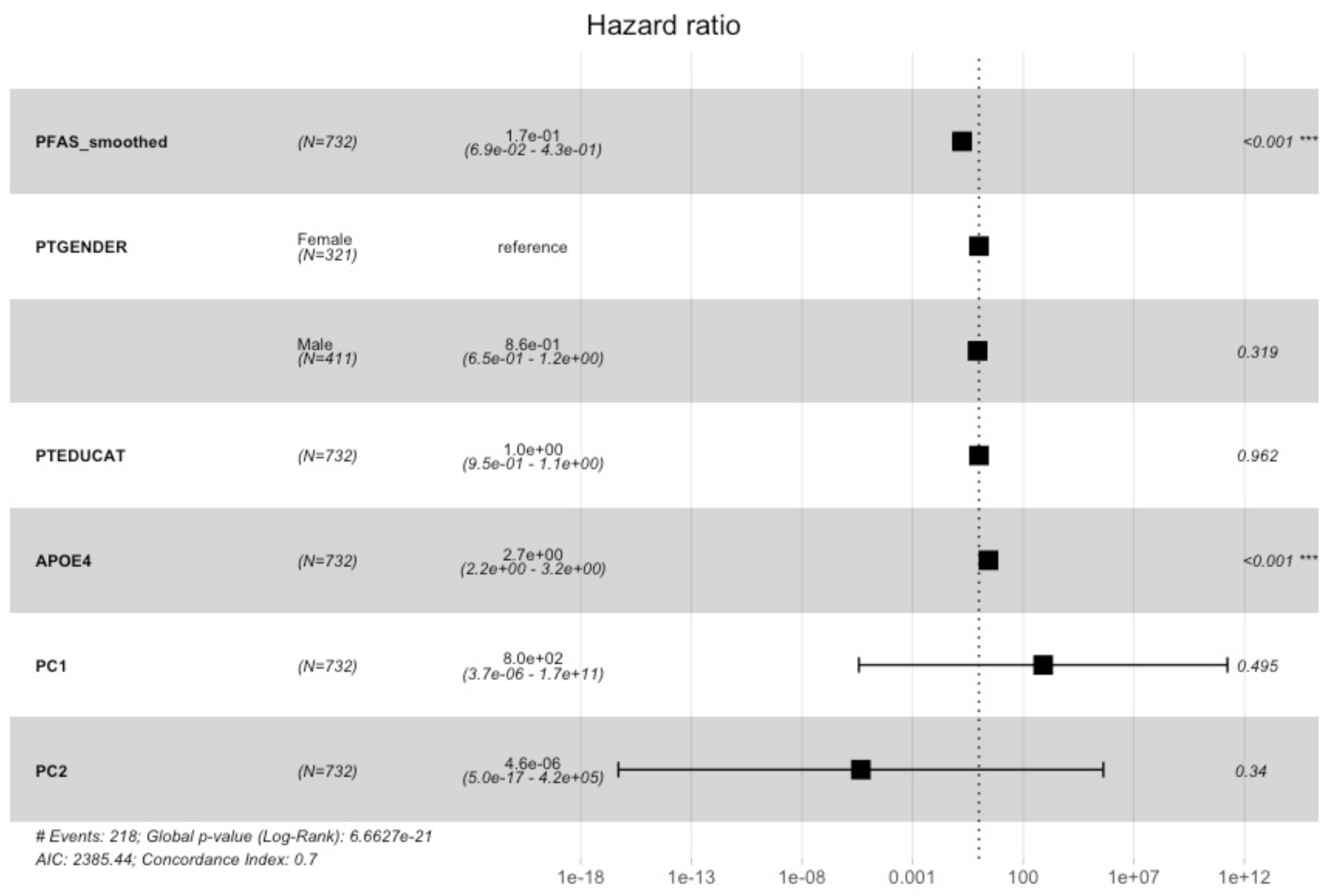
Survival analysis with the smoothed score for the gene resulting from stability selection on the ADNI dataset. The forest plot depicts hazard ratios from Cox proportional hazards model with confidence intervals and statistical significance. The score resulting from network propagation for *PFAS* was seen to be significantly associated with lower risk of conversion to AD.

### Gene set enrichment analysis

Next, we examined the community of genes surrounding *PFAS* through gene set enrichments analysis, to see if *PFAS* can be regarded as hub of a module enriched in genes related to AD. Significant overlap was found for the 1,449 genes of interest with 1,815 gene sets from Gene Ontology (GO), Chemical and Genetic Perturbations (CGP), KEGG and REACTOME, at FDR of 5%. Enrichment p-values for eight gene sets related to AD are reported in Supplementary Table 3. A high-level visual representation of 928 significantly overrepresented GO terms in their semantic space can be found in Supplementary Figure 8. The KEGG AD pathway and the four sets curated by Blalock et al. [27] comprising dysregulated genes, were all significant at FDR 5%. Moreover, randomisation control showed that the overlaps observed with the four Blalock sets could not be achieved by chance, as none of the randomly drawn gene sets returned uncorrected enrichment p-values lower than the ones observed, and only 14 out of 1,000 random sets achieved more significant overlap with the set of genes downregulated in AD curated by Wu et al. [28] (Supplementary Figure 9).

### Differential gene expression analysis

Motivated by the observed overlaps with sets of genes dysregulated in AD, we lastly sought to investigate whether the percolation of rare variants’ effects through the network impacts the transcriptomic profiles of the 30 genes identified in ADNI and ADSP, and furthermore, whether this impact exhibits tissue specificity. To test this hypothesis, we leveraged two independent, publicly available RNA-seq datasets: temporal cortex samples from the Mayo Clinic Brain Bank, and parahippocampal gyrus samples from the Mount Sinai Brain Bank.

In the Mayo RNA-seq dataset, none of the investigated genes showed significant dysregulation between controls and AD cases after correction for cell type gene markers. Full results and visualisations of the differential expression analysis in the Mount Sinai dataset for parahippocampal gyrus are reported in Supplementary Table 4 and Supplementary Figure 10. We observed a trend towards downregulation for *PFAS* and *SHOC2* in controls vs definite AD (pPFAS = 0.04; pSHOC2 = 0.001); however these did not survive multiple comparisons correction. Sixteen genes among the 30 analysed were found to be differentially expressed in at least one pairwise comparison of disease severity. A randomisation control demonstrated that this number is significantly higher than it would be expected by chance (p < 0.001; Supplementary Figure 11). In particular, significant dysregulation in controls vs definite AD was observed for *ARL1*, *ATXN10*, *CAMK2B*, *CAPNS1*, *EFNB3*, *GNB1*, *KCNMA1*, *KLC1*, *MAPRE1*, *MAPRE3*, *MOB4*, and *TREM2*.

## DISCUSSION

In this work we presented NETPAGE, a computational tool for gene-based association testing of rare variants that integrates prior knowledge about tissue-specific gene interaction networks. The aim was to boost the information content assigned to each gene, by enriching it with the knowledge of the pathways through which rare variation percolates. NETPAGE leverages the well-known strength of network propagation, combining it with multivariate sparse regression to identify genes robustly associated with a disease phenotype by analysing the genome-wide landscape as a whole and without the need for stringent statistical corrections.

NETPAGE allowed us to reveal correlations between the propagation of rare variant effects at the gene level and disease outcomes (both diagnosis and risk of conversion), as well as tissue-specific downstream effects of such propagation at the transcriptomic level. We tested the proposed method on a small dataset (ADNI), as well as on a medium-sized sample (ADSP). This enabled us to put the spotlight on a set of genes mutually connected in the hippocampus network and belonging to the same neighbourhood of diameter two. We were also able to demonstrate robustness to the choice of network by replicating the association with *PFAS* with the STRING PPI network. But one of the core features of NETPAGE is its flexibility and applicability to a wide range of traits and diseases whose genetic architecture follows the trend described by Manolio *et al.* [2].

We investigated the behaviour of the network propagation step in NETPAGE through simulated data for a range of mutation frequencies, and optimised some of the key parameters of the diffusion process. As expected, when the neighbouring genes’ mutation frequencies do not differ between cases and controls, no difference is detectable in the smoothed score of the hub gene, whereas different mutation frequencies in the neighbouring genes always flow into the hub to determine a smoothed score significantly different between cases and controls, even when the hub gene itself is not mutated, under a range of parameters and mutation scenarios. In agreement with Hofree *et al* [19], we also found the diffusion length α to have a minor, if not negligible, effect on the hub gene’s smoothed score over a sizable range and in all mutation scenarios considered. However, in contrast to Hofree et al., we found the final quantile normalisation step to be detrimental to the process and therefore excluded it from our applications to real data.

Since we regard NETPAGE as a gene-based association testing tool, we compared its performance against SKAT, a state-of-the-art method for gene- and set-based association testing of rare variants, in both datasets. In contrast to Bis *et al.* [11], SKAT-O on ADSP did not identify any significant gene, which may be due to difference in SNV inclusion criteria. Additional negative controls compared NETPAGE to mass-univariate testing of smoothed scores, stability selection on the “unsmoothed” mutation burden, and evaluated the impact of different networks (random and non-brain related). None of these techniques reported any association (Supplementary Figures 3, 5 and 6; Supplementary Table 2), clearly demonstrating the added value provided by our approach. There are many network propagation-based methods proposed over the last years, such as CATAPULT [29], HotNet2 [30], or NBS [19]. However, benchmarking NETPAGE against these methods is not a straightforward task, since these methods were designed for purposes fundamentally different from association testing, namely gene prioritisation, module detection, or patient stratification, respectively.

We also investigated the effect of different selection criteria to include SNVs in the mutation burdens to be propagated through the network. When we focused on stop-gain, stop-loss and frameshift insertions to derive our gene burdens for network propagation, we did not detect any association signal. This type of mutations has generally a higher functional impact than missense mutations, and is therefore usually subject to stronger selective pressure due to a dramatic decrease in fitness. This translates into even sparser mutation profiles, whereby the “signal enhancement” capability of network propagation is still not sufficient to reveal separation between cases and controls. We conclude that the inclusion criteria for SNVs – which are not to be regarded as a parameter of the algorithm underlying NETPAGE but rather as a user choice – need to reflect careful consideration of the disease area under study as well as the experimental conditions (sample size, sequencing technology and the like).

The genes identified by stability selection exhibit connections to a number of intermediate phenotypes and biochemical processes relevant to AD. For instance, common missense variation in *PFAS* (phosphoribosylformylglycinamidine synthase) has been linked to low-density lipoprotein (LDL) cholesterol measurement through GWAS [31,32]. Additionally, Gene Ontology annotations for *PFAS* include the term ‘glutamine metabolic process’ (GO:0006541). The metabolism of glutamine and the glutamine-glutamate cycle takes place in the brain at the astrocyte level [33]; glutamate excess in the synaptic cleft triggers neuronal excitotoxicity, which has been implicated in AD [34]. The association of rare, loss-of-function mutations in *TREM2* with AD is already well-established [35]; a discovery that historically paved the way to crucial findings about the role of neuroinflammation, microglia and innate immunity in AD [36–38]. Additionally, interactions for other NETPAGE-selected genes such as *APPBP2* and *CAPNS1* have been reported with the amyloid-precursor protein APP [39,40]. Furthermore, the local gene-interaction neighborhood of *CAMK2B*, which was identified by NETPAGE, shows a series of compelling dementia risk genes (Figure 3C): key players in familial AD (APP, PSEN2), genes in which rare variation is linked to sporadic AD (PLD3, ABCA7, SORL1; [41–43]), GWAS genes (APOE, CLU, BIN1, PICALM), genes identified through multivariate imaging genetics studies (TRIB3; [44]) and genes causal for other neurodegenerative diseases (MAPT, GRN, HTT, OTUD4 [45]). The selected hub genes therefore recapitulate the biology of some of the most important pathological processes and risk factors not only for AD, but for a broadly-defined neurodegenerative phenotype.

We demonstrated the successful application of NETPAGE to two independent datasets, one of small sample size (ADNI) and the other of moderate sample size (ADSP). In conducting these experiments, we did not seek to achieve replication of the selected genes; this task is an intrinsically challenging one, given the rarity of the mutations considered, the difference in coverage (an additional 3,000 genes were tested in ADSP that were not present in ADNI), and the small sample considered in ADNI. Our main focus was to demonstrate the validity of the proposed method in application to two very different, real-world experimental scenarios. However, it is indeed remarkable that short-range functional connections link a subset of the genes selected in the two datasets in a statistically significant fashion (Supplementary Figure 5), and we believe this to be a powerful proof of the working hypothesis underlying NETPAGE.

The role of *PFAS* as hub of a molecular pathway disrupted in AD is further strengthened by the finding of extensive and significant overlaps between the subnetwork centred on *PFAS* and the KEGG AD pathway, as well as with curated sets of genes affected by transcriptional dysregulation at distinct stages of the disease (Supplementary Table 3 and Supplementary Figure 9). Additionally, the interactome of *PFAS* appears to be enriched in genes related to a number of key ontology terms (Supplementary Figure 8), among which: biological processes such as cell ageing, microtubule-based processes, and mRNA metabolism; and cellular components such as mitochondria, myelin sheath, axon cytoplasm, and ribonucleoprotein complexes [46]. These overlaps provide additional support to the hypothesis that rare deleterious variation percolates through the network structure to significantly alter the protein landscape of the cellular environment in a pathological way. However, we also showed that these alterations are tissue-specific (and most likely even cell type-specific), since we reported differential expression for some of the core genes identified in the parahippocampal gyrus from the Mount Sinai dataset but not in samples from the broader temporal cortex in the Mayo RNA-seq dataset, at different disease stages (Supplementary Table 4). More importantly, we observed transcriptional alterations in relation to AD diagnosis in more than half of the putative core genes investigated, and demonstrated that this effect could not have been observed by chance (Supplementary Figure 11). This finding can be interpreted as pointing towards a co-expression module altered by disease. This further supports our idea that integrating information about variation and interactions is a powerful approach to gain a comprehensive, systems-level view of disease-related molecular mechanisms, as opposed to the investigation of single variants or genes of interest. However, as we did not detect differential expression for all the genes identified, we speculate that the link between these putative core genes and AD might reside in molecular mechanisms other than RNA or protein abundance, such as splicing, regulation or post-translational modifications. Some evidence in this direction is provided by a significant enrichment in genes related to post-transcriptional regulation of gene expression (Supplementary Figure 8, top panel).

We proposed NETPAGE, a methodology that enables the exploration of biological pathways through which structural variation affects tissue function and results in complex disease phenotypes. The rationale followed by NETPAGE resembles aspects of the recently proposed, although debated, omnigenic model [14,47]. There remain, however, some limitations. First, an inherent difficulty is that the ADNI dataset is a low-coverage WGS, while ADSP is WES, which can be biased due to the required exome enrichment step. Therefore, on one hand ADNI might not provide a very clean signal, while on the other hand ADSP might not enable a strict validation due to the different sequencing methods utilised. Second, we restricted this initial study design to include only rare exonic SNVs whose deleteriousness was assessed through the CADD score. Extensions of this work may address the impact on gene discovery of choices related to the study design, such as the CADD threshold for deleterious SNVs inclusion or the inclusion of intronic and regulatory variants. The use of an established sparse regression framework makes NETPAGE a highly flexible method, enabling the user to explore the relationship between smoothed mutation profiles and AD-related continuous traits, such as imaging or fluid biomarkers, beyond the simple binary definition of diagnosis.

## CONCLUSIONS

In summary, we demonstrated a novel application of network propagation to the study of rare variant effects on complex traits in two sporadic, late-onset AD cohorts. NETPAGE allowed us to identify a set of genes as tightly interconnected network hubs where the downstream influence of rare mutation accumulates and acquires predictive power for diagnosis, as well as to provide multiple lines of evidence for the biological meaning of the smoothed scores and the tissue-specific involvement of some core genes at the transcriptional level. We emphasise the flexibility of the presented methodology, that enables enhanced association testing for binary and continuous traits, as well as small and large sample sizes, as NGS is becoming an increasingly affordable alternative to SNP genotyping and adopted in many cohort and biobank studies. We believe NETPAGE is a promising approach for determining novel genetic influences on complex traits and for providing mechanistic insights into disease biology.

## MATERIALS AND METHODS

### ADNI WGS data preprocessing

Data used in the preparation of this article were obtained from the Alzheimer’s Disease Neuroimaging Initiative (ADNI) database (adni.loni.usc.edu) and the Alzheimer’s Disease Sequencing Project (ADSP) [7]. The ADNI was launched in 2003 as a public-private partnership, led by Principal Investigator Michael W. Weiner, MD. The primary goal of ADNI has been to test whether serial magnetic resonance imaging (MRI), positron emission tomography (PET), other biological markers, and clinical and neuropsychological assessment can be combined to measure the progression of mild cognitive impairment (MCI) to early AD. For up-to-date information, see www.adni-info.org. The ADNI WGS dataset was used for application to a small sample size. SNVs and small insertions-deletions (indel) data was available for 808 ADNI subjects. WGS data in Variant Call Format (VCF) was downloaded from the ADNI database. AD case or control status was available for 476 out of 808 individuals (for the remaining participants the latest diagnosis available was of MCI). Demographics and clinical outcomes for this sample are presented in Table 1.

**Table 1.** Demographics and clinical outcomes for the ADNI WGS case/control sample and the ADSP WES case/control sample.

|  | ADNI |  |  | ADSP |  |  |
| --- | --- | --- | --- | --- | --- | --- |
|  | HC | AD | Total | HC | AD | Total |
| **Sample** | 246 | 230 | 476 | 4,796 | 5,852 | 10,648 |
| **size** |  |  |  |  |  |  |
| **Women** | 134 | 92 | 226 | 2,819 | 3,371 | 6,190 |
| **White,**<br>**not**<br>**hispanic**<br>**or latino** | 222 | 217 | 439 | 4,601 | 5,585 | 10,186 |
| **Age at**<br>**latest**<br>**visit**<br>**(mean ±**<br>**sd)** | 78.3 ±<br>7.27 | 79.1 ±<br>7.58 | 78.69 ±<br>7.42 | 86.1 ± 4.53 | 75.4 ± 8.45 | 80.19 ± 8.76 |
| **APOE**<br>**ε4**<br>**(0/1/2)** | 181/64/4 | 80/117/33 | 261/181/37 | 4,080/701/15 | 3,379/2,314/159 | 7,459/3,015/174 |

### ADSP WES data preprocessing

The ADSP WES dataset (N=10,913) was then used for application to a moderate sample size. We identified 24 samples who were also sequenced as part of ADNI, who were then removed from the ADSP WES dataset, yielding a sample size of 10,889 (SI Appendix). Demographics and clinical outcomes for this final sample are presented in Table 1.

### Ancestry and population structure

Ancestry and population structure on the ADNI dataset were previously analysed from GWAS data, using SNPweights version 2.1 [48] and a two-step procedure described in [49]. We leveraged the results of this analysis to retain only study participants showing a probability of being of Caucasian ancestry greater than 80% (N = 439 AD cases and controls). For these subjects we also obtained principal components of population structure to be used in subsequent analyses.

In ADSP, due to the absence of GWAS data, individuals were filtered based on self-reported ethnicity and racial background, to include only Caucasian participants (self-reported white race and not hispanic or latino ethnicity; N = 10,413). Lastly, case-control status was available for 10,186 of these Caucasian individuals.

### Variant filtering and mapping

We used ANNOVAR, 01/06/2017 release [50] to annotate the SNVs, retaining only exonic, non-synonymous SNVs, with MAF <1% in non-Finnish-Europeans from the Exome Aggregation Consortium [51]. Variants were further filtered based on deleteriousness, by retaining SNVs ranked among the top 1% according the Combined Annotation-Dependent Depletion (CADD) method (CADD phred score>20) [52].

To assess the impact of the filtering criteria outlined above we constructed a second set of SNVs, by retaining exonic variants with MAF < 1% predicted to have a stop-gain, stop-loss or frameshift consequence. We also applied the CADD phred > 20 filter to stop-gain and stop-loss variants.

Exonic SNVs were mapped to genes according to annotations based on the RefSeq database [53]. These filtering and mapping procedures were applied to the ADNI and ADSP datasets separately.

### Tissue-specific gene interaction networks

As a substrate for network propagation, we leveraged tissue-specific weighted gene interaction networks from Greene et al. [15]. In these networks, each node represents a gene, each edge a functional relationship, and an edge between two genes is probabilistically weighted based on experimental evidence connecting both genes. We focused on the interaction network for the human hippocampus, being the key brain structure related to the loss of episodic memory in AD [54]. Additional negative controls were performed using: a randomised version of the same hippocampus network, obtained by shuffling edges after binarisation (see next section); and a gene network not related to brain tissue, specifically the umbilical cord network.

Lastly, we investigated the robustness of the method with respect to the network structure by using the human, non-tissue-specific protein-protein interaction network (PPI) available through the STRING database [21].

### Propagation of mutation signals through gene interaction networks

To model the propagation of the effects of rare mutations through a gene interaction network, we adopted the *network propagation* approach first introduced by [55] for semi-supervised learning. In our case, a gene carrying a deleterious variant is used as a seed in an iterative procedure that propagates its effect according to the network structure. Hence the effect propagation effectively reproduces a graph-constrained diffusion process whereby information from the seed gene flows not only to its first neighbours but to all genes in a connected component, and the amount of information flowing into a gene is determined by the strength of its connections to the source gene.

Practically, a single individual’s whole genome data is represented as a gene-based vector of length M, where for each gene the mutational burden can be encoded as either the count of rare deleterious variants (rare-variant burden) or a binary variable indicating the presence or absence of any such mutation (rare-variant status). Stacking these vectors of all S subjects yields the sparse matrix *G^0^* ∈ ℕ*^S^*^x*M*^. A network can be either in the form of a weighted graph, represented by a square, symmetric, similarity matrix *N* ∈ ℝ*^M^*^x*M*^ with *0* ≤ *N_ij_* ≤ *1*, or in the form of an unweighted graph, represented by a square, symmetric adjacency matrix *N* ∈ ℕ*^M^*^x*M*^ with *N_ij_* ∈ {*0*; *1*}. The gene burden *G^0^* is propagated through the network simultaneously for all subjects according to the following iterative procedure:

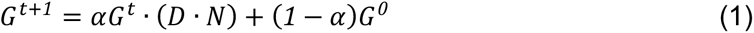

where *G^t^*^+1^ is the smoothed mutation profile at iteration *t* + *1*; *α* ∈ [0,1] is a tuning parameter governing the distance a signal is allowed to diffuse through the network, and *D* ∈ ℝ*^M^*^x*M*^ is a diagonal matrix with the inverse of the node strengths of *N* along the diagonal. Equation (1) is iteratively evaluated until convergence (i.e., until the L^2^-norm of *G^t^*^+1^ – *G^t^* is smaller than a pre-specified threshold).

Our implementation of network propagation in NETPAGE allows the user to specify: whether the network used is a gene- or a protein-interaction network, and the gene/protein naming convention; the type of encoding used in the input file (i.e., if rare-variant burden or rare-variant status is used); the convergence threshold on ‖*G^t^*^+1^ – *G^t^*‖*_2_* (default 10^-6^); the diffusion length α (default 0.5); this was optimized through simulation; whether the full weighted graph is to be used to guide the diffusion process, or if a graph adjacency matrix is to be generated from the original graph by retaining only the top P% edges (we refer to this procedure as network binarisation); the percentage P of top edges to be retained, in case network binarisation is to be performed (default 1%); this was optimized through simulation; whether the gene interaction network features self-loops (default False); whether the rows of the smoothed mutation profile are to be quantile-normalised after convergence (default False). Quantile normalisation is the last step performed after network propagation by NBS [19], to ensure that the smoothed mutation profile for each patient follows the same distribution.

Network propagation is sensitive to the direction of the mutation’s effect (i.e., protective vs. deleterious), therefore the user can choose if and how to deal with the direction of effects of the genes’ mutational burdens with respect to a given binary phenotype. Briefly, the bioinformatics assessment of deleteriousness (e.g., CADD score) is unrelated to any disease phenotype, hence the mutational status of a given gene can be equally risk-increasing or protective with respect to a specific binary phenotype, resulting in blended effects in signal-receiving genes. Therefore, we require a mechanism to numerically distinguish the “signal” flowing into a hub from a protective gene from the “signal” coming from a risk gene. In light of this, we are providing the user with additional flexibility to either: set to 0 the mutation status/burden of protective genes; set to 0 the mutation status/burden of risk genes; set the mutation status to −1 for protective genes and to +1 for risk genes; none of the above. Risk and protective genes are determined by the direction of effect of their mutation status on the case-control status (odds ratio from a Fisher’s test on the 2×2 contingency table).

### NETPAGE: application to AD sequencing data

We performed network propagation on the full (i.e., without filtering on ancestry or diagnosis) ADNI and ADSP datasets separately. Each subject-level mutational burden (row of the *G^0^*matrix) was encoded as a binary vector of length M (0 = gene not carrying any of the selected SNVs, 1 = gene carrying at least one of the selected SNVs). The *G^0^* matrix in ADNI described the mutation status of 13,310 genes for 808 study participants. The *G^0^*matrix in ADSP described the mutation status of 16,268 genes for 10,889 study participants. In both applications, we used the default values for the parameters α and P, as optimised through simulations, and set the mutation status to −1 for protective genes and to +1 for risk genes.

### Stability selection

In order to identify genes robustly associated with disease, smoothed mutation profiles resulting from network propagation were related to clinical diagnosis via sparse logistic regression. We applied LASSO regression [22] as implemented in the R package *glmnet* [56]. Briefly, *glmnet* finds the coefficients’ vector β that solves the following regularised regression problem:

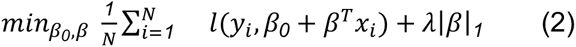

over a range of values for λ. Here *l*(*y*;, *y*;*_est_*) is the negative log-likelihood contribution for observation *i*. In Equation (2), *N* is the number of observations (subjects), *β_0_* is the model intercept and *β* the vector of regression coefficients. For subject *i*, *y*;*_i_* is the value of the response variable, and *x_i_* a vector of predictors. The LASSO or L1-regularisation corresponds to minimising the L1-norm of the coefficients’ vector |*β*|*_1_*. As a consequence, most of the coefficients in *β* shrink to zero, achieving efficient variable selection. The amount of regularisation is controlled by the *λ* parameter. Here, we performed stability selection [23], as implemented in the R package *stabs* [57], to estimate the optimal amount of regularisation required. In brief, stability selection combines variable selection with bootstrap resampling to estimate a probability value for each variable to be selected by the sparse regression. For a given regression task, we performed 100 bootstrap (split-half) resamplings and focused on smoothed gene scores with a selection probability higher than 80%. Stability selection in ADNI was performed on smoothed scores for M = 13,310 genes and N = 439 Caucasian subjects (222 healthy controls [HC], 217 AD; date accessed April 24th, 2018). Stability selection in ADSP was performed on smoothed scores for 16,298 genes and 10,186 Caucasian subjects (4601 HC, 5585 AD).

### Gene-based rare variant association testing

We conducted gene-based association testing for rare, deleterious SNVs with clinical diagnosis in ADNI with a state-of-the-art method, in order to benchmark NETPAGE’s performance. Specifically, we used SKAT-O [25], which combines burden and variance-component tests, implemented in the R package *SKAT* [10]. Diagnosis at the latest available time point was coded as a dichotomous trait (HC vs AD). Associations were tested for M = 13,591 genes controlling for age, sex, number of *APOE* ε4 alleles, years of education, and two principal components of CEU population substructure (see section *Ancestry and population structure*). Genome-wide significance was established at p < 0.05/13,591 = 3.6×10^-6^. Additionally, to demonstrate the advantage of our multivariate approach, a mass univariate test of association between the smoothed gene scores and case-control status was also conducted via logistic regression for M = 13,310 genes, controlling for the same confounders listed above. Genome-wide significance was established at p < 0.05/13,310 = 3.75×10^-6^. The sample for both these tests comprised 439 Caucasian cases and controls from ADNI.

We further conducted a gene-based association test with SKAT-O on the same 10,186 European subjects from ADSP used for stability selection; 270,165 non-synonymous SNVs were grouped into 16,630 genes. Associations were tested correcting for age, sex and number of APOE4 alleles. The aim of this test was to benchmark NETPAGE against a state-of-the-art method on a dataset that enables sufficient statistical power to detect associations, as it has been already reported [11].

### Set-based rare variant association testing

We conducted set-based association testing for rare, deleterious SNVs with case/control status in ADNI. Gene sets were defined from the results of stability selection: each selected gene was grouped with its first neighbours in the hippocampus network. Burden, variance-component, and omnibus (SKAT-O) tests were performed with this set definition.

### Model comparison

After stability selection, we focused on the smoothed score of the top selected gene. With it, we built two logistic regression models: a baseline model, including case-control status as response, and as predictors the same set of covariates used for SKAT; and an extended model, adding to the predictors in the baseline model the smoothed score of the selected gene. Sample size for these models was 439 Caucasian cases and controls from ADNI. We compared the goodness-of-fit of the two models through a chi-squared test. Additionally, we computed the pseudo-R^2^ statistic (the analogue of the percentage of variance explained in linear regression models) for the baseline and the extended model. We used the McFadden method in the PseudoR2 function in the R package *DescTools* [58]. We repeated the same procedure for the ADSP-selected genes, one at a time. P-values from the chi-squared tests were corrected for multiple comparisons using the Bonferroni procedure (Table 2). The predictors for the baseline model now included only sex, age and number of *APOE* ε4 alleles. Years of education was not recorded among the ADSP phenotypes. It was also not possible to compute CEU population substructure, due to lack of GWAS data. Sample size for these models was 10,186 Caucasian cases and controls.

**Table 2.** Selection probabilities for the 30 genes whose smoothed score was identified as robust predictor of case-control status in ADNI and ADSP. Genes highlighted in bold are second or closer neighbours of *PFAS* in the hippocampus network.

| Gene | Selection probability | Model comparison p-value (Bonferroni corrected) | Gene | Selection probability | Model comparison p-value (Bonferroni corrected) |
| --- | --- | --- | --- | --- | --- |
| **NETPAGE - ADNI** | | | ***GNB1***<br>chr 1p36.33 | 1.000 | $1 \times 10^{-29}$ |
| *PFAS*<br>chr 17p13.1 | 0.850 | $2 \times 10^{-4}$ | ***HIC2***<br>chr 22q11.21 | 0.990 | $1 \times 10^{-11}$ |
| **NETPAGE - ADSP** | | | ***KCNMA1***<br>chr 10q22.3 | 0.980 | $4 \times 10^{-13}$ |
| ***ABR***<br>chr 17p13.3 | 0.845 | $4 \times 10^{-16}$ | ***KLC1***<br>chr 14q32.33 | 1.000 | $1 \times 10^{-23}$ |
| ***ADRM1***<br>chr 20q13.33 | 0.965 | $6 \times 10^{-16}$ | ***MAPK11***<br>chr 22q13.33 | 0.990 | $1 \times 10^{-18}$ |
| *APPBP2*<br>chr 17q23.2 | 0.905 | $4 \times 10^{-22}$ | ***MAPRE1***<br>chr 20q11.21 | 0.825 | $1 \times 10^{-16}$ |
| ***ARL1***<br>chr 12q23.2 | 0.990 | $3 \times 10^{-22}$ | ***MAPRE3***<br>chr 2p23.3 | 0.970 | $1 \times 10^{-13}$ |
| *ATXN10*<br>chr 22q13.31 | 0.965 | $7 \times 10^{-17}$ | ***MOB4***<br>chr 5p13.1-p12 | 0.895 | $1 \times 10^{-28}$ |
| *CAMK2B*<br>chr 7p13 | 0.960 | $1 \times 10^{-7}$ | ***MRPL17***<br>chr 11p15.4 | 0.910 | $1 \times 10^{-24}$ |
| **CAPNS1**<br>chr 19q13.12 | 0.990 | 5x10<sup>-19</sup> | **PPP1CC**<br>chr 12q24.11 | 0.965 | 1x10<sup>-36</sup> |
| **COPS5**<br>chr 8q13.1 | 0.935 | 4x10<sup>-24</sup> | **RAB1A**<br>chr 2p14 | 0.845 | 2x10<sup>-12</sup> |
| **CSNK1A1**<br>chr 5q32 | 1.000 | 1x10<sup>-36</sup> | **SHOC2**<br>chr 10q25.2 | 0.995 | 1x10<sup>-28</sup> |
| **CUL5**<br>chr 11q22.3 | 0.995 | 4x10<sup>-27</sup> | **TMEM147**<br>chr 19q13.12 | 0.985 | 1x10<sup>-24</sup> |
| **DCTN6**<br>chr 8p12 | 0.940 | 2x10<sup>-27</sup> | **TREM2**<br>chr 6p21.1 | 0.990 | 0.14 |
| **DSTN**<br>chr 20p12.1 | 0.935 | 2x10<sup>-33</sup> | **UBL3**<br>chr 13q12.3 | 1.000 | 4x10<sup>-33</sup> |
| **EFNB3**<br>chr 17p13.1 | 0.820 | 2x10<sup>-4</sup> | **ZNF207**<br>chr 17q11.2 | 0.995 | 3x10<sup>-27</sup> |

### Survival analysis

We conducted survival analysis on the selected genes in ADNI with the R packages *survival* [59] and *survminer* [60]. The event considered was either conversion from HC to AD or conversion from MCI to AD. The time variable was age at dementia onset if conversion occurred; or age at latest diagnosis if conversion did not occur (right-censored time-to-event). Sample size was N = 732 ADNI Caucasian subjects with WGS data (>80% CEU ancestry). For the selected genes we considered both the “raw” (0/1) mutation status, and the smoothed score derived with network propagation. For each gene, we first modelled and compared survival curves for mutation carriers vs non-carriers (log-rank test); we then refined the comparison of mutation carriers and non-carriers by fitting Cox proportional hazards models, controlling for sex, number of *APOE* ε4 alleles, years of education, and two principal components of population. Lastly, we fitted the same Cox models replacing the binary gene mutation status with its smoothed score, retaining the same set of covariates. Because healthy controls were already used for stability selection, in order to mitigate the risk of a circular analysis we repeated the analysis excluding stable healthy controls (NHC = 222, final sample size N = 510), therefore focusing only on stable MCI who did not convert to AD as the event-free population.

### Gene-set enrichment analysis

Gene set enrichment analysis was performed on the gene selected in the ADNI dataset, together with its first and second neighbours in the thresholded and binarised hippocampus gene network. This yielded a set of 1,449 genes to be tested. As background set, we used the set of M = 13,310 genes resulting from network propagation on the ADNI dataset. We tested for enrichment using the Gene Ontology (GO) and chemical and genetic perturbation (CGP) gene sets, as well as the KEGG and REACTOME pathway collections available in the Molecular Signatures Database v6.1 [61]. Overrepresentation of the selected genes in these curated sets was tested with the Fisher’s exact test. Enrichment p-values were adjusted for multiple comparisons using the Benjamini-Hochberg procedure at a false discovery rate of 5%. Significant GO terms were aggregated in hierarchies and visualised through ReViGO (http://revigo.irb.hr/revigo.jsp) [62]. For seven CGP gene sets relevant to Alzheimer’s disease, we further applied a randomisation procedure to ensure that the observed enrichments could not be achieved by chance. We formed 1,000 set replicates by randomly drawing 1,449 genes from the background; every set replicate was tested for overrepresentation in the seven CGP sets, then the p-values from the original set were compared to the distributions of p-values from the randomised sets.

### Differential gene expression analysis

Motivated by the observed overlaps with sets of genes dysregulated in AD, we lastly sought to investigate whether differential expression between cases and controls occurs for genes selected in ADNI and ADSP. In order to investigate also the tissue-specificity of this effect, we leveraged two independent RNA-seq datasets: the Mayo Clinic Brain Bank (MCBB) [63] and the Mount Sinai Brain Bank (MSBB) [64]. Within these datasets, we specifically focused on post-mortem expression levels in the temporal cortex and parahippocampal gyrus (Brodmann area 36), respectively. This choice aimed at maximising proximity and consistency with the hippocampal gene interaction network used in the discovery phase, being aware of the high degree of spatial variability in transcriptomics patterns across the brain. All data was accessed through the AMP-AD Knowledge Portal (www.synapse.org). For the Mayo dataset, we leveraged publicly available results of differential expression analysis, with and without correction for neuronal marker gene levels (Synapse ID syn6090802). RNA sequencing and processing at the MSBB was described in detail elsewhere [64]. Each sample was assigned a neuropathology category according to the Consortium to Establish a Registry for Alzheimer’s Disease (CERAD) protocol (1=normal, 2=definite AD, 3=probable AD, 4=possible AD) [65]. We formed a gene set of interest comprising genes selected in ADNI and ADSP; we then used the Dunn’s test for stochastic dominance [66] to perform pairwise nonparametric testing for differences in normalised gene expression levels between CERAD neuropathology categories in this gene set of interest. P-values were corrected for gene-wise, pairwise comparisons using the Benjamini-Hochberg procedure at a false discovery rate of 5%, and the Bonferroni method for the number of genes tested. Lastly, we performed a randomisation control to quantify the probability of seeing N genes with a significant differential expression among M genes randomly drawn from the MSBB dataset. We formed 1000 sets of M randomly drawn genes, conducted the Dunn’s test for each gene and set, and counted how many genes in each random set showed significantly different expression in at least one pairwise comparison. We then plotted the distribution of the number of randomly dysregulated genes and compared it to the actual number of dysregulated genes found in the original set of interest.

## DECLARATIONS

### Ethics approval and consent to participate

Statement on ethics approval and consent for ADNI and ADSP can be found in the Supplementary Materials.

### Consent for publication

Not applicable

### Availability of data and materials

Accession codes for the datasets used in this study can be found in the Supplementary Materials. A Python 2.7 module for network propagation will be released upon publication at https://github.com/maffleur/NETPAGE.git.

### Authors’ contributions

MAS and AA designed the study; VN and MDG provided data; MAS analysed data; MAS, VN and AA wrote manuscript; all authors reviewed and approved manuscript.

## Acknowledgments

Some of the computing for this project was performed on the Sherlock 2.0 cluster. We would like to thank Stanford University and the Stanford Research Computing Center for providing computational resources and support that contributed to these research results. We would also like to thank Prof Jonathan M. Schott from the UCL Dementia Research Centre for his valuable clinical insights.

## References

1. Locke AE, Kahali B, Berndt SI, Justice AE, Pers TH, Day FR, et al. Genetic studies of body mass index yield new insights for obesity biology. Nature. 2015;

2. Manolio TA, Collins FS, Cox NJ, Goldstein DB, Hindorff LA, Hunter DJ, et al. Finding the missing heritability of complex diseases. Nature. 2009.

3. Saunders a M, Strittmatter WJ, Schmechel D, George-Hyslop PH, Pericak-Vance M a, Joo SH, et al. Association of apolipoprotein E allele epsilon 4 with late-onset familial and sporadic Alzheimer’s disease. Neurology. 1993;43:1467–72.

4. Corder EH, Saunders AM, Strittmatter WJ, Schmechel DE, Gaskell PC, Small GW, et al. Gene Dose of Apolipoprotein-E Type-4 Allele and the Risk of Alzheimers-Disease in Late-Onset Families. Science (80-.). 1993;261:921–3.

5. Cruchaga C, Chakraverty S, Mayo K, Vallania FLM, Mitra RD, Faber K, et al. Rare variants in APP, PSEN1 and PSEN2 increase risk for AD in late-onset Alzheimer’s disease families. PLoS One. 2012;

6. Rohrer JD, Warren JD. Phenotypic signatures of genetic frontotemporal dementia. Curr. Opin. Neurol. 2011.

7. Beecham GW, Bis JC, Martin ER, Choi S-H, DeStefano AL, van Duijn CM, et al. The Alzheimer’s Disease Sequencing Project: Study design and sample selection. Neurol. Genet. 2017;

8. Lee S, Abecasis GR, Boehnke M, Lin X. Rare-Variant Association Analysis: Study Designs and Statistical Tests. Am. J. Hum. Genet. [Internet]. 2014 [cited 2017 Sep 22];95:5–23. Available from: https://ac.els-cdn.com/S0002929714002717/1-s2.0-S0002929714002717-main.pdf?_tid=b3ad90c2-9f86-11e7-9ffe-00000aab0f27&acdnat=1506078834_2681f860cb507432912772ead161bb2a

9. Cohen JC, Kiss RS, Pertsemlidis A, Marcel YL, McPherson R, Hobbs HH. Multiple rare alleles contribute to low plasma levels of HDL cholesterol. Science (80-.). 2004;

10. Wu MC, Lee S, Cai T, Li Y, Boehnke M, Lin X. Rare-variant association testing for sequencing data with the sequence kernel association test. Am. J. Hum. Genet. 2011;

11. Bis JC, Jian X, Kunkle BW, Chen Y, Hamilton-Nelson KL, Bush WS, et al. Whole exome sequencing study identifies novel rare and common Alzheimer’s-Associated variants involved in immune response and transcriptional regulation. Mol. Psychiatry. Nature Publishing Group; 2018;

12. Purcell SM, Moran JL, Fromer M, Ruderfer D, Solovieff N, Roussos P, et al. A polygenic burden of rare disruptive mutations in schizophrenia. Nature. 2014;

13. Walter K, Min JL, Huang J, Crooks L, Memari Y, McCarthy S, et al. The UK10K project identifies rare variants in health and disease. Nature. 2015;

14. Boyle EA, Li YI, Pritchard JK. An Expanded View of Complex Traits: From Polygenic to Omnigenic [Internet]. Cell. 2017 [cited 2017 Jun 19]. p. 1177–86. Available from: http://dx.doi.org/10.1016/j.cell.2017.05.038

15. Greene CS, Krishnan A, Wong AK, Ricciotti E, Zelaya RA, Himmelstein DS, et al. Understanding multicellular function and disease with human tissue-specific networks. Nat. Genet. [Internet]. 2015;47:569–76. Available from: http://www.nature.com/doifinder/10.1038/ng.3259%Cnhttp://www.ncbi.nlm.nih.gov/pubmed/25915600%Cnhttp://www.pubmedcentral.nih.gov/articlerender.fcgi?artid=PMC4828725

16. Cowen L, Ideker T, Raphael BJ, Sharan R. Network propagation: A universal amplifier of genetic associations. Nat. Rev. Genet. [Internet]. Nature Publishing Group; 2017;18:551–62. Available from: http://dx.doi.org/10.1038/nrg.2017.38

17. Vanunu O, Magger O, Ruppin E, Shlomi T, Sharan R. Associating genes and protein complexes with disease via network propagation. PLoS Comput. Biol. 2010;6.

18. Winter C, Kristiansen G, Kersting S, Roy J, Aust D, Knösel T, et al. Google goes cancer: Improving outcome prediction for cancer patients by network-based ranking of marker genes. PLoS Comput. Biol. 2012;8.

19. Hofree M, Shen JP, Carter H, Gross A, Ideker T. Network-based stratification of tumor mutations. Nat. Methods [Internet]. Nature Research; 2013 [cited 2017 Feb 20];10:1108–15. Available from: http://www.nature.com/doifinder/10.1038/nmeth.2651

20. Magger O, Waldman YY, Ruppin E, Sharan R. Enhancing the Prioritization of Disease-Causing Genes through Tissue Specific Protein Interaction Networks. PLoS Comput. Biol. 2012;

21. Szklarczyk D, Franceschini A, Wyder S, Forslund K, Heller D, Huerta-Cepas J, et al. STRING v10: protein-protein interaction networks, integrated over the tree of life. Nucleic Acids Res. [Internet]. 2015 [cited 2017 May 8];43:D447–52. Available from: http://www.ncbi.nlm.nih.gov/pubmed/25352553

22. Hastie T, Tibshirani R, Friedman J. The Elements of Statistical Learning: Data Mining, Inference, and Prediction, Second Edition (Springer Series in Statistics) (9780387848570): Trevor Hastie, Robert Tibshirani, Jerome Friedman: Books. Elem. Stat. Learn. dta mining, inference, Predict. 2011.

23. Meinshausen N, Bühlmann P. Stability selection. J. R. Stat. Soc. Ser. B Stat. Methodol. 2010;

24. Kunkle BW, Grenier-Boley B, Sims R, Bis JC, Damotte V, Naj AC, et al. Genetic meta-analysis of diagnosed Alzheimer’s disease identifies new risk loci and implicates Aβ, tau, immunity and lipid processing. Nat. Genet. [Internet]. Nature Publishing Group; 2019 [cited 2019 Mar 4];51:414–30. Available from: http://www.nature.com/articles/s41588-019-0358-2

25. Lee S, Emond MJ, Bamshad MJ, Barnes KC, Rieder MJ, Nickerson DA, et al. Optimal unified approach for rare-variant association testing with application to small-sample case-control whole-exome sequencing studies. Am. J. Hum. Genet. 2012;

26. Richardson TG, Timpson NJ, Campbell C, Gaunt TR. A pathway-centric approach to rare variant association analysis. Eur. J. Hum. Genet. [Internet]. Nature Publishing Group; 2017 [cited 2019 Jan 29];25:123–9. Available from: http://www.nature.com/articles/ejhg2016113

27. Blalock EM, Geddes JW, Chen KC, Porter NM, Markesbery WR, Landfield PW. Incipient Alzheimer’s disease: microarray correlation analyses reveal major transcriptional and tumor suppressor responses. Proc. Natl. Acad. Sci. U. S. A. [Internet]. 2004 [cited 2019 Mar 4];101:2173–8. Available from: http://www.pnas.org/lookup/doi/10.1073/pnas.0308512100

28. Wu Z, Guo H, Chow N, Sallstrom J, Bell RD, Deane R, et al. Role of the MEOX2 homeobox gene in neurovascular dysfunction in Alzheimer disease. Nat. Med. [Internet]. 2005 [cited 2019 Mar 4];11:959–65. Available from: http://www.nature.com/articles/nm1287

29. Singh-Blom UM, Natarajan N, Tewari A, Woods JO, Dhillon IS, Marcotte EM. Prediction and Validation of Gene-Disease Associations Using Methods Inspired by Social Network Analyses. Aloy P, editor. PLoS One [Internet]. Public Library of Science; 2013 [cited 2019 Mar 4];8:e58977. Available from: http://dx.plos.org/10.1371/journal.pone.0058977

30. Leiserson MDM, Vandin F, Wu H-T, Dobson JR, Eldridge J V, Thomas JL, et al. Pan-cancer network analysis identifies combinations of rare somatic mutations across pathways and protein complexes. Nat. Genet. [Internet]. Nature Publishing Group; 2015 [cited 2019 Mar 4];47:106–14. Available from: http://www.nature.com/articles/ng.3168

31. Below JE, Parra EJ, Gamazon ER, Torres J, Krithika S, Candille S, et al. Meta-analysis of lipid-traits in Hispanics identifies novel loci, population-specific effects, and tissue-specific enrichment of eQTLs. Sci. Rep. [Internet]. 2016 [cited 2019 Mar 4];6:19429. Available from: http://www.nature.com/articles/srep19429

32. Spracklen CN, Chen P, Kim YJ, Wang X, Cai H, Li S, et al. Association analyses of East Asian individuals and trans-ancestry analyses with European individuals reveal new loci associated with cholesterol and triglyceride levels. Hum. Mol. Genet. [Internet]. 2017 [cited 2019 Mar 4];26:1770–84. Available from: https://academic.oup.com/hmg/article/26/9/1770/3039197

33. Schousboe A, Scafidi S, Bak LK, Waagepetersen HS, McKenna MC. Glutamate Metabolism in the Brain Focusing on Astrocytes. Adv. Neurobiol. [Internet]. 2014 [cited 2019 Mar 4]. p. 13–30. Available from: http://www.ncbi.nlm.nih.gov/pubmed/25236722

34. Hynd MR, Scott HL, Dodd PR. Glutamate-mediated excitotoxicity and neurodegeneration in Alzheimer’s disease. Neurochem. Int. [Internet]. Pergamon; 2004 [cited 2019 Mar 4];45:583–95. Available from: https://www.sciencedirect.com/science/article/pii/S0197018604000555

35. Guerreiro R, Wojtas A, Bras J, Carrasquillo M, Rogaeva E, Majounie E, et al. TREM2 variants in Alzheimer’s disease. N. Engl. J. Med. [Internet]. 2013;368:117–27. Available from: http://www.nejm.org/doi/abs/10.1056/NEJMoa1211851%5Cnhttp://www.ncbi.nlm.nih.gov/pubmed/23150934%5Cnhttp://www.pubmedcentral.nih.gov/articlerender.fcgi?artid=PMC3631573

36. Ulrich JD, Ulland TK, Colonna M, Holtzman DM. Elucidating the Role of TREM2 in Alzheimer’s Disease. Neuron [Internet]. Elsevier; 2017 [cited 2019 Mar 4];94:237–48. Available from: http://www.ncbi.nlm.nih.gov/pubmed/28426958

37. Ulland TK, Song WM, Huang SC-C, Ulrich JD, Sergushichev A, Beatty WL, et al. TREM2 Maintains Microglial Metabolic Fitness in Alzheimer’s Disease. Cell [Internet]. Elsevier; 2017 [cited 2019 Mar 4];170:649–663.e13. Available from: http://www.ncbi.nlm.nih.gov/pubmed/28802038

38. Li J-T, Zhang Y. TREM2 regulates innate immunity in Alzheimer’s disease. J. Neuroinflammation [Internet]. BioMed Central; 2018 [cited 2019 Mar 4];15:107. Available from: https://jneuroinflammation.biomedcentral.com/articles/10.1186/s12974-018-1148-y

39. Zheng P, Eastman J, Vande Pol S, Pimplikar SW. PAT1, a microtubule-interacting protein, recognizes the basolateral sorting signal of amyloid precursor protein. Proc. Natl. Acad. Sci. U. S. A. [Internet]. 1998 [cited 2019 Mar 4];95:14745–50. Available from: http://www.ncbi.nlm.nih.gov/pubmed/9843960

40. Lee M, Kwon YT, Li M, Peng J, Friedlander RM, Tsai L-H. Neurotoxicity induces cleavage of p35 to p25 by calpain. Nature [Internet]. 2000 [cited 2019 Mar 4];405:360–4. Available from: http://www.ncbi.nlm.nih.gov/pubmed/10830966

41. Steinberg S, Stefansson H, Jonsson T, Johannsdottir H, Ingason A, Helgason H, et al. Loss-of-function variants in ABCA7 confer risk of Alzheimer’s disease. Nat. Genet. 2015;

42. Cruchaga C, Karch CM, Jin SC, Benitez BA, Cai Y, Guerreiro R, et al. Rare coding variants in the phospholipase D3 gene confer risk for Alzheimer’s disease. Nature. 2014;

43. Vardarajan BN, Zhang Y, Lee JH, Cheng R, Bohm C, Ghani M, et al. Coding mutations in SORL1 and Alzheimer disease. Ann. Neurol. 2015;

44. Lorenzi M, Altmann A, Gutman B, Wray S, Arber C, Hibar DP, et al. Susceptibility of brain atrophy to TRIB3 in Alzheimer’s disease, evidence from functional prioritization in imaging genetics. Proc. Natl. Acad. Sci. 2018;201706100.

45. Margolin DH, Kousi M, Chan Y-M, Lim ET, Schmahmann JD, Hadjivassiliou M, et al. Ataxia, Dementia, and Hypogonadotropism Caused by Disordered Ubiquitination. From Dep. Neurol. N Engl J Med [Internet]. 2013 [cited 2017 Dec 12];21368:1992–2003. Available from: http://www.nejm.org/doi/pdf/10.1056/NEJMoa1215993

46. Bai B, Wu H, Street C, Hanfelt J, Cheng D, Jin P, et al. U1 small nuclear ribonucleoprotein complex and RNA splicing alterations in Alzheimer’s disease. Proc. Natl. Acad. Sci. 2013;

47. Wray NR, Wijmenga C, Sullivan PF, Yang J, Visscher PM. Common Disease Is More Complex Than Implied by the Core Gene Omnigenic Model. Cell [Internet]. 2018 [cited 2018 Aug 21];173:1573–80. Available from: https://doi.org/10.1016/j.cell.2018.05.051

48. Chen CY, Pollack S, Hunter DJ, Hirschhorn JN, Kraft P, Price AL. Improved ancestry inference using weights from external reference panels. Bioinformatics [Internet]. Oxford University Press; 2013 [cited 2016 Dec 19];29:1399–406. Available from: http://www.ncbi.nlm.nih.gov/pubmed/23539302

49. Scelsi MA, Khan RR, Lorenzi M, Christopher L, Greicius MD, Schott JM, et al. Genetic study of multimodal imaging Alzheimer’s disease progression score implicates novel loci. Brain. 2018;141:2167–80.

50. Wang K, Li M, Hakonarson H. ANNOVAR: Functional annotation of genetic variants from high-throughput sequencing data. Nucleic Acids Res. 2010;38.

51. Lek M, Karczewski KJ, Minikel E V., Samocha KE, Banks E, Fennell T, et al. Analysis of protein-coding genetic variation in 60,706 humans. Nature [Internet]. Nature Research; 2016 [cited 2016 Aug 23];536:285–91. Available from: http://www.nature.com/doifinder/10.1038/nature19057

52. Kircher M, Witten DM, Jain P, O BJ, Cooper GM, Shendure J. A general framework for estimating the relative pathogenicity of human genetic variants. Nat. Publ. Gr. [Internet]. 2014 [cited 2017 Jun 19];46. Available from: https://www.nature.com/ng/journal/v46/n3/pdf/ng.2892.pdf

53. Pruitt KD, Tatusova T, Maglott DR. NCBI reference sequences (RefSeq): A curated non-redundant sequence database of genomes, transcripts and proteins. Nucleic Acids Res. 2007;

54. West MJ, Coleman PD, Flood DG, Troncoso JC. Differences in the pattern of hippocampal neuronal loss in normal ageing and Alzheimer’s disease. Lancet. 1994;

55. Zhou D, Bousquet O, Lal TN, Weston J, Schölkopf B. Learning with local and global consistency. Adv. neural … [Internet]. 2004;1:595–602. Available from: http://citeseerx.ist.psu.edu/viewdoc/summary?doi=10.1.1.115.3219%5Cnhttp://books.google.com/books?hl=en&lr=&id=0F-9C7K8fQ8C&oi=fnd&pg=PA321&dq=Learning+with+Local+and+Global+Consistency&ots=TGLsqTUb52&sig=PtUN7Swm8rO0fEFlPtGEhYZ00MA

56. Friedman AJ, Hastie T, Simon N, Tibshirani R, Hastie MT. Lasso and Elastic-Net Regularized Generalized Linear Models. Available online https://cran.r-project.org/web/packages/glmnet/glmnet.pdf. (Verified 29 July. 2015). 2015.

57. Hofner B, Boccuto L, Göker M. Controlling false discoveries in high-dimensional situations: Boosting with stability selection. BMC Bioinformatics. 2015;

58. Signorell A. DescTools: Tools for descriptive statistics. R package version 0.99.20. CRAN. 2017;

59. Therneau T. Package Survival: A Package for Survival Analysis in R. R Packag. version 2.38. 2015;

60. Kassambara A, Kosinski M, Biecek P, Fabian S. survminer: Drawing Survival Curves using “ggplot2”. R package. version 0.4.3. 2018.

61. Subramanian A, Tamayo P, Mootha VK, Mukherjee S, Ebert BL, Gillette MA, et al. Gene set enrichment analysis: a knowledge-based approach for interpreting genome-wide expression profiles. Proc. Natl. Acad. Sci. U. S. A. [Internet]. National Academy of Sciences; 2005 [cited 2017 Nov 28];102:15545–50. Available from: http://www.ncbi.nlm.nih.gov/pubmed/16199517

62. Supek F, Bošnjak M, Škunca N, Šmuc T. Revigo summarizes and visualizes long lists of gene ontology terms. PLoS One. 2011;

63. Allen M, Carrasquillo MM, Funk C, Heavner BD, Zou F, Younkin CS, et al. Human whole genome genotype and transcriptome data for Alzheimer’s and other neurodegenerative diseases. Sci. Data [Internet]. Nature Publishing Group; 2016 [cited 2019 Mar 4];3:160089. Available from: http://www.nature.com/articles/sdata201689

64. Wang M, Beckmann ND, Roussos P, Wang E, Zhou X, Wang Q, et al. The Mount Sinai cohort of large-scale genomic, transcriptomic and proteomic data in Alzheimer’s disease. Sci. Data [Internet]. Nature Publishing Group; 2018 [cited 2019 Mar 4];5:180185. Available from: http://www.nature.com/articles/sdata2018185

65. Mirra SS, Heyman A, McKeel D, Sumi SM, Crain BJ, Brownlee LM, et al. The Consortium to Establish a Registry for Alzheimer’s Disease (CERAD). Part II. Standardization of the neuropathologic assessment of Alzheimer’s disease. Neurology [Internet]. 1991 [cited 2019 Mar 4];41:479–86. Available from: http://www.ncbi.nlm.nih.gov/pubmed/2011243

66. Dunn OJ. Multiple Comparisons Using Rank Sums. Technometrics [Internet]. 1964 [cited 2019 Mar 4];6:241–52. Available from: http://www.tandfonline.com/doi/abs/10.1080/00401706.1964.10490181

