## Supplementary Materials for "Network propagation of rare mutations in Alzheimer’s disease reveals tissue-specific hub genes and communities"

**This PDF file includes:**

Supplementary text
Tables S1 to S4
Figs. S1 to S11
References for SI reference citations

**Supplementary Information Text**

#### Genetic data processing

*ADNI WGS data preprocessing*

WGS was obtained from blood genomic DNA samples and processed by ADNI, as described elsewhere (1). Briefly, sequencing was performed using the Illumina HiSeq2000 system through paired-end read chemistry and read lengths of 100 base pairs. Reads were aligned to the reference human genome (NCBI build 37.72) using the BWA tool (2) and used for multi-sample variant calling with the GATK HaplotypeCaller (3).

*ADSP WES data preprocessing*

ADSP WES data were retrieved through dbGAP (accession ID: phs000572.v7.p4), which include QC'ed SNV genotypes concordant between the Atlas (Baylor's) and GATK (Broad's) calling pipelines. Briefly, 10,913 subjects underwent WES at three different sequencing centers (the Human Genome Sequencing Center at Baylor College of Medicine, Broad Institute, and Genome Institute at Washington University), capturing the exome target region using the Illumina Rapid Capture Exome or the Nimblegen's VCRome v2.1 exome kits and paired-end sequencing them on Illumina HiSeq 2000 platform. Detailed information about sequencing and QC pipeline is available from https://www.niagads.org/adsp/content/sequencing-pipelines. Whole-genome sequencing (WGS) data, available from ADSP was not considered since the majority of the subjects available were of Caribbean-Hispanic ancestry.

We identified ADSP subjects who were sequenced as part of ADNI and removed them from the sample. To identify overlapping samples, we performed the following steps: we selected from the ADNI WGS data the set of exonic SNVs in common with ADSP WES, then merged the two datasets using PLINK v1.9 (4). On the merged dataset, we performed basic quality control (minor allele frequency [MAF] <0.05, SNV missingness rate >0.1, Hardy-Weinberg equilibrium p<5e-7), followed by linkage disequilibrium-based pruning (PLINK parameters: --indep-pairwise 500 50 0.2). On this dataset of independent exonic SNVs, we computed identity-by-descent for all pairs of samples, and identified as duplicates all ADSP subjects exhibiting PI_HAT > 0.95 (PLINK parameters: --genome --min 0.95).

#### Tissue-specific gene interaction networks

As a substrate for network propagation, we leveraged tissue-specific weighted gene interaction networks from Greene et al. (5). In these networks, each node represents a gene, each edge a functional relationship, and an edge between two genes is probabilistically weighted based on experimental evidence connecting both genes. Curation of these networks involved the integration of evidence from 987 genome-scale data sets encompassing approximately 38,000 conditions from an estimated 14,000 publications including both expression and interaction measurements in 144 tissues and cell lineages, with each data set weighted in a process specific manner. Tissue-specific gene interaction networks are freely available for download from http://hb.flatironinstitute.org/download.

**Network propagation - synthetic data**
We conducted a set of experiments with simulated gene mutation profiles to examine the behaviour of our network propagation implementation, as well as to determine the optimal values for some of the parameters listed in the previous section.
Simulation and investigation of the parameter space was carried out jointly in three nested levels. Further details can be found in Online Methods.
***Level 1*:** the original, weighted network is read in; the network is then binarised retaining a certain percentage P of top edges. We vary $P \in \{0.5; 1; 5; 10; 25\}$. For the sake of computational speed, we do not focus on the entire hippocampus gene network; instead, from the binarised network we extract a subgraph of radius 2 centered on a randomly selected hub gene (in this instance, *CALML3*), using the python package *networkx* version 2.2 (6). The number M of nodes in this subgraph is retained for use in the next level.
***Level 2*:** synthetic, binary mutation profiles for S = 2,000 subjects (equally divided into AD cases and controls) and M genes are generated by assigning the "mutated" status to proportions $f_{HC}$ and $f_{AD}$ of controls and cases randomly for each gene. We vary the gene-level mutation frequencies in the following ranges:

- mutation frequency in controls $f_{HC} \in \{0; 0.1; 0.5\}\%$
- mutation frequency in AD cases $f_{AD}\{0.1; 0.5; 1\}\%$

Additionally, we simulate three different scenarios for the propagation of mutation signals:

1. only the first neighbours of the hub gene are mutated;
2. only the second neighbours of the hub gene are mutated;
3. both first and second neighbours are mutated (realistic scenario).

This results in 27 synthetic mutation profiles generated. This is repeated also allowing the hub gene to be mutated.
***Level 3:*** for each of the 27 previously generated synthetic mutation profiles, network propagation is run by varying:

- the diffusion length $\alpha \in \{0; 0.25; 0.5; 0.75; 0.9\}$;
- whether quantile normalisation is applied or not to the final smooth profile. Quantile normalisation was used in Hofree et al. (7) for clustering purposes; however, we sought to assess its impact for the purpose of association testing.

Lastly, after network propagation was run with the selected set of parameters on the simulated data, the smoothed score for the hub gene was tested for difference between cases and controls with a Wilcoxon rank-sum test. Statistical significance was established at $p < 0.05$.

**Differential gene expression analysis**
RNA sequencing and processing at the Mayo Clinic Brain Bank was described in detail elsewhere ((8) and https://www.synapse.org/#!Synapse:syn3163039 ). Each sample was assigned one of the following pathological diagnoses: Alzheimer's disease (AD, N=84), progressive supranuclear palsy (PSP, N=84), pathologic aging (PA, N=30), and control (HC, N=80). We only focused on differential expression analysis for HC vs AD (total N = 156 after sample QC). Normalized read counts were assessed for differential expression between diagnosis groups, using multi-variable linear regression adjusting for key

covariates. Two models were run for each comparison called “Simple” (syn6090804) and “Comprehensive” (syn6090803). The simple model includes as covariates: age at death, sex, RNA integrity number (RIN), Source and FLOWCELL (syn3817650). The comprehensive model includes the same covariates plus normalized counts for 5 genes as surrogate variables for relevant cell types as follows: *CD68* (Microglia), *CD34* (Endothelial cells), *OLIG2* (Oligodendroglia), *GFAP* (Astrocytes) and *ENO2* (Neurons).

RNA sequencing and processing at the Mount Sinai Brain Bank was described in detail elsewhere ((9) and https://www.synapse.org/#!Synapse:syn3157743). Gene expression levels were normalised by regressing out the effect of sex, race, age at death, post-mortem interval, RIN, exonic rate, rRNA rate and batch. The accession code for normalised expression levels in Brodmann area 36 is syn16795937. We also downloaded RNA-seq covariates (syn6100548) and post-mortem clinical assessments (syn6101474). Each sample was assigned a neuropathology category according to the Consortium to Establish a Registry for Alzheimer's Disease (CERAD) protocol (1=normal, 2=definite AD, 3=probable AD, 4=possible AD) (10).

**Data availability and funding**

Data collection and sharing for this project was funded by the Alzheimer's Disease Neuroimaging Initiative (ADNI) (National Institutes of Health Grant U01 AG024904) and DOD ADNI (Department of Defense award number W81XWH-12-2-0012). ADNI is funded by the National Institute on Aging, the National Institute of Biomedical Imaging and Bioengineering, and through generous contributions from the following: AbbVie, Alzheimer's Association; Alzheimer's Drug Discovery Foundation; Araclon Biotech; BioClinica, Inc.; Biogen; Bristol-Myers Squibb Company; CereSpir, Inc.; Cogstate; Eisai Inc.; Elan Pharmaceuticals, Inc.; Eli Lilly and Company; EuroImmun; F. Hoffmann-La Roche Ltd and its affiliated company Genentech, Inc.; Fujirebio; GE Healthcare; IXICO Ltd.; Janssen Alzheimer Immunotherapy Research \& Development, LLC.; Johnson \& Johnson Pharmaceutical Research \& Development LLC.; Lumosity; Lundbeck; Merck \& Co., Inc.; Meso Scale Diagnostics, LLC.; NeuroRx Research; Neurotrack Technologies; Novartis Pharmaceuticals Corporation; Pfizer Inc.; Piramal Imaging; Servier; Takeda Pharmaceutical Company; and Transition Therapeutics. The Canadian Institutes of Health Research is providing funds to support ADNI clinical sites in Canada. Private sector contributions are facilitated by the Foundation for the National Institutes of Health (www.fnih.org). The grantee organization is the Northern California Institute for Research and Education, and the study is coordinated by the Alzheimer’s Therapeutic Research Institute at the University of Southern California. ADNI data are disseminated by the Laboratory for Neuro Imaging at the University of Southern California.

The Alzheimer’s Disease Sequencing Project (ADSP) is comprised of two Alzheimer’s Disease (AD) genetics consortia and three National Human Genome Research Institute (NHGRI) funded Large Scale Sequencing and Analysis Centers (LSAC). The two AD genetics consortia are the Alzheimer’s Disease Genetics Consortium (ADGC) funded by NIA (U01 AG032984), and the Cohorts for Heart and Aging Research in Genomic Epidemiology (CHARGE) funded by NIA (R01 AG033193), the National Heart, Lung, and Blood Institute (NHLBI), other National Institute of Health (NIH) institutes and other foreign governmental and non-governmental organizations. The Discovery Phase analysis of sequence data is supported through UF1AG047133 (to Drs. Schellenberg, Farrer, Pericak-Vance, Mayeux, and Haines); U01AG049505 to Dr. Seshadri; U01AG049506 to Dr. Boerwinkle; U01AG049507 to Dr. Wijsman; and U01AG049508 to Dr. Goate and the Discovery Extension Phase analysis is supported through U01AG052411 to Dr. Goate, U01AG052410 to Dr. Pericak-Vance and U01 AG052409 to Drs. Seshadri and Fornage. Data generation and harmonization in the Follow-up Phases is supported by U54AG052427 (to Drs. Schellenberg and Wang).

The ADGC cohorts include: Adult Changes in Thought (ACT), the Alzheimer's Disease Centers (ADC), the Chicago Health and Aging Project (CHAP), the Memory and Aging Project (MAP), Mayo Clinic (MAYO), Mayo Parkinson's Disease controls, University of Miami, the Multi-Institutional Research in Alzheimer's Genetic Epidemiology Study (MIRAGE), the National Cell Repository for Alzheimer's Disease (NCRAD), the National Institute on Aging Late Onset Alzheimer's Disease Family Study (NIA-LOAD), the Religious Orders Study (ROS), the Texas Alzheimer's Research and Care Consortium (TARC), Vanderbilt University/Case Western Reserve University (VAN/CWRU), the Washington Heights-Inwood Columbia Aging Project (WHICAP) and the Washington University Sequencing Project (WUSP), the Columbia University Hispanic- Estudio Familiar de Influencia Genetica de Alzheimer (EFIGA), the University of Toronto (UT), and Genetic Differences (GD).

The CHARGE cohorts are supported in part by National Heart, Lung, and Blood Institute (NHLBI) infrastructure grant HL105756 (Psaty), RC2HL102419 (Boerwinkle) and the neurology working group is supported by the National Institute on Aging (NIA) R01 grant AG033193. The CHARGE cohorts participating in the ADSP include the following: Austrian Stroke Prevention Study (ASPS), ASPS-Family study, and the Prospective Dementia Registry-Austria (ASPS/PRODEM-Aus), the Atherosclerosis Risk in Communities (ARIC) Study, the Cardiovascular Health Study (CHS), the Erasmus Rucphen Family Study (ERF), the Framingham Heart Study (FHS), and the Rotterdam Study (RS). ASPS is funded by the Austrian Science Fond (FWF) grant number P20545-P05 and P13180 and the Medical University of Graz. The ASPS-Fam is funded by the Austrian Science Fund (FWF) project I904),the EU Joint Programme - Neurodegenerative Disease Research (JPND) in frame of the BRIDGET project (Austria, Ministry of Science) and the Medical University of Graz and the Steiermärkische Krankenanstalten Gesellschaft. PRODEM-Austria is supported by the Austrian Research Promotion agency (FFG) (Project No. 827462) and by the Austrian National Bank (Anniversary Fund, project 15435. ARIC research is carried out as a collaborative study supported by NHLBI contracts (HHSN268201100005C, HHSN268201100006C, HHSN268201100007C, HHSN268201100008C, HHSN268201100009C, HHSN268201100010C, HHSN268201100011C, and HHSN268201100012C). Neurocognitive data in ARIC is collected by U01 2U01HL096812, 2U01HL096814, 2U01HL096899, 2U01HL096902, 2U01HL096917 from the NIH (NHLBI, NINDS, NIA and NIDCD), and with previous brain MRI examinations funded by R01-HL70825 from the NHLBI. CHS research was supported by contracts HHSN268201200036C, HHSN268200800007C, N01HC55222, N01HC85079, N01HC85080, N01HC85081, N01HC85082, N01HC85083, N01HC85086, and grants U01HL080295 and U01HL130114 from the NHLBI with additional contribution from the National Institute of Neurological Disorders and Stroke (NINDS). Additional support was provided by R01AG023629, R01AG15928, and R01AG20098 from the NIA. FHS research is supported by NHLBI contracts N01-HC-25195 and HHSN268201500001I. This study was also supported by additional grants from the NIA (R01s AG054076, AG049607 and AG033040 and NINDS (R01 NS017950). The ERF study as a part of EUROSPAN (European Special Populations Research Network) was supported by European Commission FP6 STRP grant number 018947 (LSHG-CT-2006-01947) and also received funding from the European Community's Seventh Framework Programme (FP7/2007-2013)/grant agreement HEALTH-F4-2007-201413 by the European Commission under the programme "Quality of Life and Management of the Living Resources" of 5th Framework Programme (no. QLG2-CT-2002-01254). High-throughput analysis of the ERF data was supported by a joint grant from the Netherlands Organization for Scientific Research and the Russian Foundation for Basic Research (NWO-RFBR 047.017.043). The Rotterdam Study is funded by Erasmus Medical Center and Erasmus University, Rotterdam, the Netherlands Organization for Health Research and Development (ZonMw), the Research Institute for Diseases in the Elderly (RIDE), the Ministry of Education, Culture and Science, the Ministry for Health, Welfare and Sports, the European Commission (DG XII), and the municipality of Rotterdam. Genetic data sets are also supported by the Netherlands Organization of Scientific Research NWO Investments (175.010.2005.011, 911-03-012), the Genetic Laboratory of the Department of Internal Medicine, Erasmus MC, the Research Institute for Diseases in the Elderly (014-93-015; RIDE2), and the Netherlands Genomics Initiative (NGI)/Netherlands Organization for Scientific Research (NWO) Netherlands Consortium for Healthy Aging (NCHA), project 050-060-810. All studies are grateful to their participants, faculty and staff. The content of these manuscripts is solely the responsibility of the authors and does not necessarily represent the official views of the National Institutes of Health or the U.S. Department of Health and Human Services.

The four LSACs are: the Human Genome Sequencing Center at the Baylor College of Medicine (U54 HG003273), the Broad Institute Genome Center (U54HG003067), The American Genome Center at the Uniformed Services University of the Health Sciences (U01AG057659), and the Washington University Genome Institute (U54HG003079).

Biological samples and associated phenotypic data used in primary data analyses were stored at Study Investigators institutions, and at the National Cell Repository for Alzheimer’s Disease (NCRAD, U24AG021886) at Indiana University funded by NIA. Associated Phenotypic Data used in primary and secondary data analyses were provided by Study Investigators, the NIA funded Alzheimer’s Disease Centers (ADCs), and the National Alzheimer’s Coordinating Center (NACC, U01AG016976) and the National Institute on Aging Genetics of Alzheimer’s Disease Data Storage Site (NIAGADS, U24AG041689) at the University of Pennsylvania, funded by NIA, and at the Database for

Genotypes and Phenotypes (dbGaP) funded by NIH. This research was supported in part by the Intramural Research Program of the National Institutes of health, National Library of Medicine. Contributors to the Genetic Analysis Data included Study Investigators on projects that were individually funded by NIA, and other NIH institutes, and by private U.S. organizations, or foreign governmental or nongovernmental organizations.

Study data were provided by the following sources: The Mayo Clinic Alzheimers Disease Genetic Studies, led by Dr. Nilufer Taner and Dr. Steven G. Younkin, Mayo Clinic, Jacksonville, FL using samples from the Mayo Clinic Study of Aging, the Mayo Clinic Alzheimers Disease Research Center, and the Mayo Clinic Brain Bank. Data collection was supported through funding by NIA grants P50 AG016574, R01 AG032990, U01 AG046139, R01 AG018023, U01 AG006576, U01 AG006786, R01 AG025711, R01 AG017216, R01 AG003949, NINDS grant R01 NS080820, CurePSP Foundation, and support from Mayo Foundation. Study data includes samples collected through the Sun Health Research Institute Brain and Body Donation Program of Sun City, Arizona. The Brain and Body Donation Program is supported by the National Institute of Neurological Disorders and Stroke (U24 NS072026 National Brain and Tissue Resource for Parkinsons Disease and Related Disorders), the National Institute on Aging (P30 AG19610 Arizona Alzheimers Disease Core Center), the Arizona Department of Health Services (contract 211002, Arizona Alzheimers Research Center), the Arizona Biomedical Research Commission (contracts 4001, 0011, 05-901 and 1001 to the Arizona Parkinson's Disease Consortium) and the Michael J. Fox Foundation for Parkinsons Research.

**Supplementary Tables**

**Table S1** – Selection probabilities (from the application of NETPAGE to ADSP) for the 21 genes reported in the recent GWAS by [1]**.**

| Gene | Selection probability | Gene | Selection probability |
|---|---|---|---|
| *CR1* | 0.00 | *FERMT2* | 0.00 |
| *BIN1* | 0.00 | *SLC24A4* | 0.00 |
| *INPP5D* | NA | *ABCA7* | 0.00 |
| *HLA-DRB1* | 0.00 | *APOE* | 0.00 |
| *TREM2* | 0.99 | *CASS4* | 0.00 |
| *CD2AP* | 0.00 | *ECHDC3* | 0.00 |
| *NYAP1* | 0.00 | *ACE* | 0.00 |
| *EPHA1* | 0.00 | *NDUFAF6* | 0.00 |
| *PTK2B* | 0.00 | *ADAM10* | 0.02 |
| *CLU* | 0.00 | *IQCK* | 0.00 |
| *SPI1* | 0.00 | *MIR142* | NA |
| *MS4A2* | 0.00 | *ADAMTS1* | 0.00 |
| *PICALM* | 0.18 | *OARD1* | 0.00 |
| *SORL1* | 0.00 | *WWOX* | 0.00 |

**Table S2 -** Results of set-based SKAT test for association of rare, exonic, deleterious variants with case-control status in ADNI. The set of interest was formed by including the first interaction neighbours of PFAS. Additionally, the first two rows in the table report gene-based p-values for PFAS from Supplementary Figure 5.

| **Test** | **PFAS p-value** |
|---|---|
| Omnibus (SKAT-O), gene-based | 0.39 |
| Smoothed logistic, gene-based | 0.001 |
| Burden, set-based | 0.82 |
| Variance-component, set-based | 0.43 |
| Omnibus (SKAT-O), set-based | 0.66 |

**Table S3 -** Results of gene set enrichment analysis for eight AD-related gene sets. The gene set tested for overrepresentation included PFAS and its first and second interaction neighbours in the thresholded and binarised hippocampus functional network. Hits = number of observed genes overlapping with the curated gene set of interest; expected = number of genes expected to overlap with the curated gene set of interest by chance; OR = odds ratio from the Fisher's test; P = p-value of the Fisher's test; $P_{corr}$ = p-value corrected with the Benjamini-Hochberg procedure. P-values reported in bold are significant at FDR 5%.

| **Gene set** | **Hits** | **Expected** | **OR** | **P** | **$P_{corr}$ (FDR)** |
|---|---|---|---|---|---|
| Alzheimer's disease (KEGG) | 22 | 9.68 | 2.70 | 0.00016 | **0.00145** |
| Blalock - upregulated in incipient AD (CGP) | 242 | 127.47 | 2.35 | 6.18E-25 | **4.18E-23** |
| Blalock - downregulated in AD (CGP) | 214 | 75.25 | 4.12 | 9.39E-50 | **2.33E-47** |
| Blalock - upregulated in AD (CGP) | 227 | 178.01 | 1.38 | 0.00003 | **0.00077** |
| Blalock - downregulated in incipient AD (CGP) | 33 | 10.87 | 4.09 | 2.35E-9 | **4.25E-8** |
| Ray - Alzheimer's disease (CGP) | 0 | 0.86 | 0 | 1 | 1 |
| Wu - upregulated in AD (CGP) | 0 | 0.86 | 0 | 1 | 1 |
| Wu - downregulated in AD (CGP) | 3 | 1.07 | 3.55 | 0.08392 | 0.32 |

**Table S4** - Differential expression analysis uncorrected, two-tailed p-values on the MSBB RNA-seq dataset for the 30 genes selected from ADNI and ADSP. The + or – sign in brackets next to the p-value represents the direction of effect detected. Pairwise comparisons were performed for normalised expression levels against CERAD diagnosis using the nonparametric Dunn's test. Values in boldface are the ones that remained significant after a two-fold multiple testing correction (Benjamini-Hochberg for the multiple pairwise comparisons for a given gene, Bonferroni for the number of genes tested). In total, 16 genes were seen to be significantly dysregulated in at least one comparison.

| Gene | Normal vs definite AD | Normal vs probable AD | Normal vs possible AD | Definite vs probable AD | Definite vs possible AD | Probable vs possible AD |
|---|---|---|---|---|---|---|
| *ABR* | 0.08532549 (+) | 0.43979832 (+) | 0.19530774 (+) | 0.55995801 (-) | 0.9375899 (+) | 0.6012055 (+) |
| *ADRM1* | 0.06367708 (+) | 0.07145437 (+) | 0.53333357 (-) | 0.70854412 (+) | 0.04665654 (-) | 0.04490362 (-) |
| *APPBP2* | 0.02484203 (+) | 0.20420772 (+) | 0.31279853 (-) | 0.6246178 (-) | 0.00759026 (-) | 0.05516895 (-) |
| *ARL1* | **9.60E-05 (+)** | 0.06868582 (+) | 0.52088796 (-) | 0.2113508 (-) | **0.00046906 (-)** | 0.04166424 (-) |
| *ATXN10* | **2.80E-05 (+)** | 0.04086053 (+) | 0.35591872 (-) | 0.21149131 (-) | **6.43E-05 (-)** | 0.01363344 (-) |
| *CAMK2B* | **0.00016811 (+)** | 0.150141 (+) | 0.27336492 (+) | 0.12517956 (-) | 0.1067733 (-) | 0.84791951 (-) |
| *CAPNS1* | **0.00025439 (+)** | 0.07254326 (+) | 0.62529398 (-) | 0.27980563 (-) | 0.00155892 (-) | 0.05998178 (-) |
| *COPS5* | 0.00710867 (+) | 0.20371741 (+) | 0.09812357 (-) | 0.39679236 (-) | **0.00025426 (-)** | 0.01280683 (-) |
| *CSNK1A1* | 0.22103889 (+) | 0.17192355 (+) | 0.00941451 (-) | 0.66790396 (+) | 0.00037365 (-) | 0.00067422 (-) |
| *CUL5* | 0.05242147 (+) | 0.30949872 (+) | 0.1487663 (-) | 0.61220644 (-) | 0.00379395 (-) | 0.03593812 (-) |
| *DCTN6* | 0.0027226 (+) | 0.11422272 (+) | 0.47348518 (-) | 0.43888473 (-) | 0.00352703 (-) | 0.05630661 (-) |

| | | | | | | |
|---|---|---|---|---|---|---|
| *DSTN* | 0.00042003 (+) | 0.08444402 (+) | 0.35031492 (-) | 0.29442766 (-) | **0.00042157 (-)** | 0.02651501 (-) |

| | | | | | | |
|---|---|---|---|---|---|---|
| *EFNB3* | **9.13E-05 (+)** | 0.64852753 (+) | 0.90548686 (-) | 0.00759608 (-) | 0.00298583 (-) | 0.63634429 (-) |
| *GNB1* | **2.27E-06 (+)** | 0.01168865 (+) | 0.51526791 (+) | 0.23412477 (-) | 0.00557148 (-) | 0.14469109 (-) |
| *HIC2* | 0.4273242 (-) | 0.15210139 (-) | 0.33693627 (+) | 0.39954048 (-) | 0.11779922 (+) | 0.0449449 (+) |
| *KCNMA1* | **7.86E-05 (+)** | 0.00076011 (+) | 0.248018 (+) | 0.75733279 (+) | 0.09164148 (-) | 0.09021318 (-) |
| *KLC1* | **0.00016008 (+)** | 0.74519054 (+) | 0.83310809 (-) | 0.00705741 (-) | 0.00304198 (-) | 0.6536745 (-) |
| *MAPK11* | 0.00090225 (+) | 0.29087258 (+) | 0.10484374 (+) | 0.11570227 (-) | 0.45359383 (-) | 0.55992478 (+) |
| *MAPRE1* | **3.48E-05 (-)** | 0.42846987 (-) | 0.56359731 (-) | 0.0122629 (+) | 0.01556105 (+) | 0.89829388 (+) |
| *MAPRE3* | **2.25E-05 (+)** | 0.11549068 (+) | 0.46637302 (+) | 0.07575642 (-) | 0.01948768 (-) | 0.5317311 (-) |
| *MOB4* | **0.00010529 (+)** | 0.01812549 (+) | 0.36298807 (-) | 0.50195552 (-) | **0.00017224 (-)** | 0.00669804 (-) |
| *MRPL17* | 0.0008427 (+) | 0.70798635 (+) | 0.02820632 (-) | 0.02187392 (-) | **2.82E-06 (-)** | 0.02483284 (-) |
| *PFAS* | 0.04544683 (+) | 0.71003601 (-) | 0.4337333 (+) | 0.04661365 (-) | 0.51466858 (-) | 0.32086484 (+) |
| *PPP1CC* | 0.05580053 (-) | 0.56580786 (-) | 0.67483367 (-) | 0.34625704 (+) | 0.33663905 (+) | 0.92664189 (+) |
| *RAB1A* | 0.00212783 (+) | 0.10804656 (+) | 0.17125365 (-) | 0.42064694 (-) | **0.00027171 (-)** | 0.01216714 (-) |

| | | | | | | |
|---|---|---|---|---|---|---|
| *SHOC2* | 0.00184187 (+) | 0.04086053 (+) | 0.52276831 (-) | 0.6985566 (-) | 0.00346631 (-) | 0.02673288 (-) |
| *TMEM147* | 0.06657185 (+) | 0.61425714 (+) | 0.15208069 (-) | 0.34120129 (-) | 0.0050111 (-) | 0.09413337 (-) |
| *TREM2* | **4.52E-06 (-)** | 0.26407975 (-) | 0.05616947 (-) | 0.01150605 (+) | 0.16905671 (+) | 0.43021011 (-) |
| *UBL3* | 0.7909294 (+) | 0.75982655 (+) | 0.34345886 (-) | 0.9179262 (+) | 0.24356415 (-) | 0.27794552 (-) |
| *ZNF207* | 0.05803408 (-) | 0.67046254 (-) | 0.13017915 (-) | 0.27936106 (+) | 0.8607827 (-) | 0.31931312 (-) |

**Figure S1** - A selection of simulation results. The x and y axis represent the mutation frequencies in controls and cases respectively. The faceting allows to visualise the effect of other parameters (diffusion length, percentage of edges retained, quantile normalisation). The colour-coding indicates the statistical significance of the difference in the hub gene smoothed score between controls and cases, in units of -log10 p-value. We investigated three mutation scenarios: scenario 1, only first neighbours mutated; scenario 2, only second neighbours mutated; scenario 3, both first and second neighbours mutated; the target gene is always unmutated. Cells marked with a black cross indicate parameter combinations where the smooth score of the hub gene was not significantly different between cases and controls. (A) Effect of quantile normalisation after network propagation with top 1% edges retained (top row, scenario 1 on the left, scenario 3 on the right), and with top 25% edges retained (bottom row, mutation scenarios as in top row). (B) Joint effect of percentage of edges retained and diffusion length for the three mutation scenarios considered.

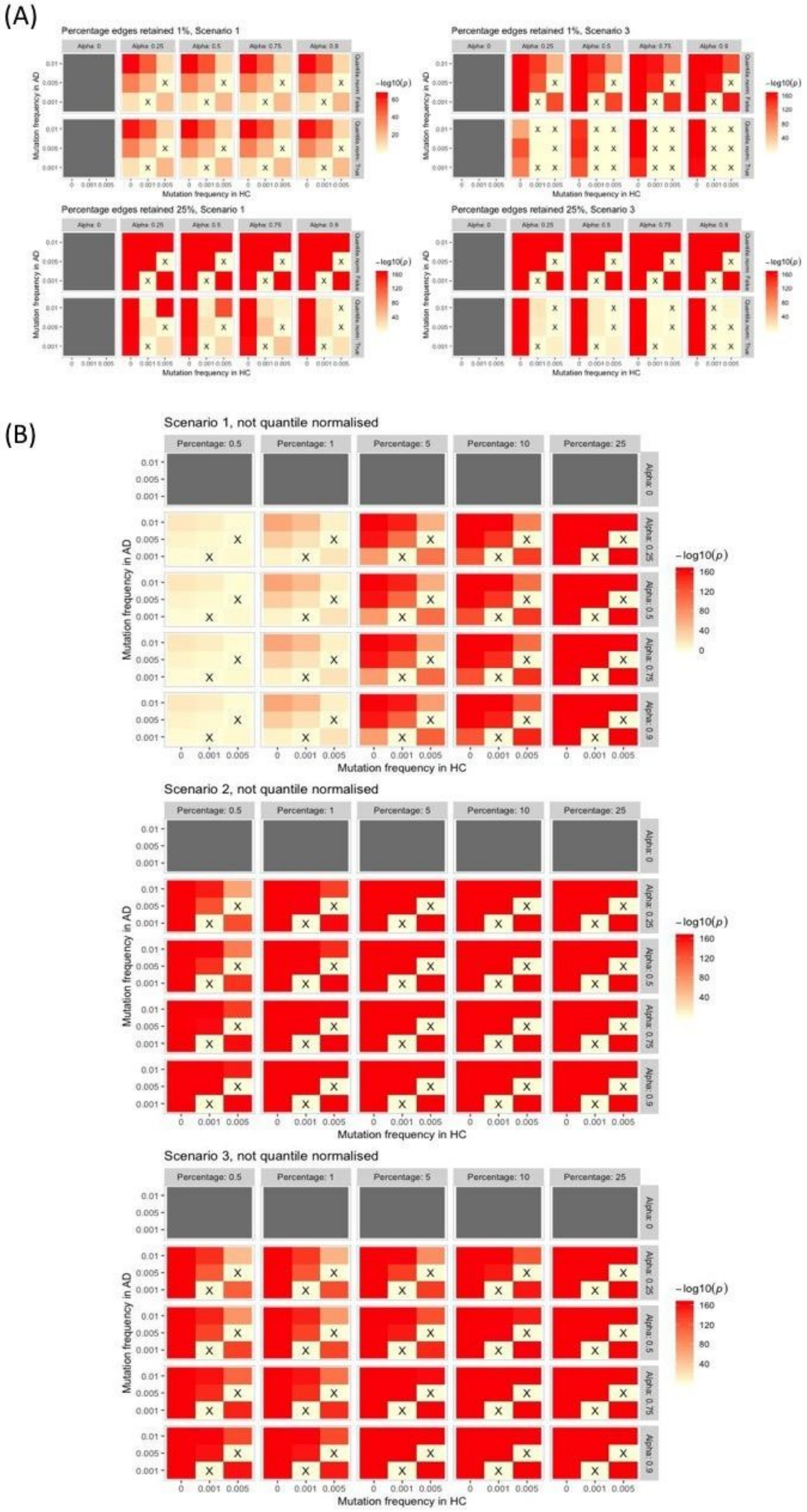
(A)
Percentage edges retained 1%, Scenario 1
Percentage edges retained 1%, Scenario 3
Percentage edges retained 25%, Scenario 1
Percentage edges retained 25%, Scenario 3
(B)
Scenario 1, not quantile normalised
Scenario 2, not quantile normalised
Scenario 3, not quantile normalised
Percentage: 0.5
Percentage: 1
Percentage: 5
Percentage: 10
Percentage: 25
Alpha: 0
Alpha: 0.25
Alpha: 0.5
Alpha: 0.75
Alpha: 0.9
Mutation frequency in AD
Mutation frequency in HC
0.01
0.005
0.001
0
-log10(p)
160
120
80
40

**Figure S2 -** Stability selection results in ADNI for a selection probability cutoff of 0.80, after propagating the burden of rare variants through the STRING PPI network. In these stability paths plots, the x axis represents the steps taken along the lambda sequence to choose the optimal amount of regularisation required by the LASSO; the y axis represents the selection probability of a gene. Selection probability paths (trajectories) for different genes are represented by different colours. Genes whose trajectories crossed the threshold of 0.80 selection probability were considered as robust predictors of case-control status and followed up in subsequent analyses. The selection of *PFAS* is here replicated.

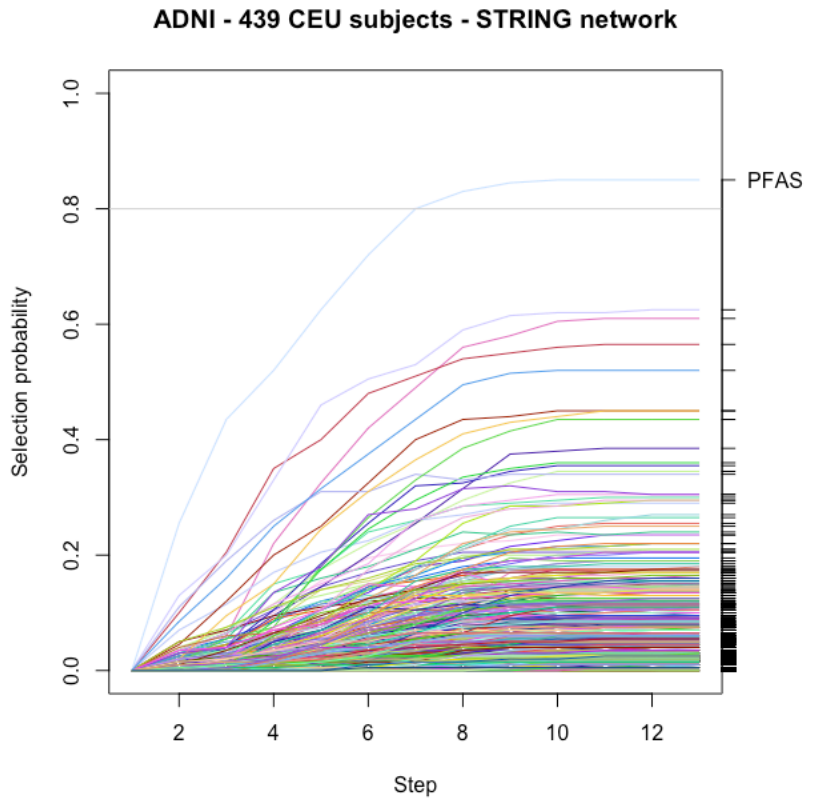

**Figure S3** - Stability selection on (A) the raw ("unsmoothed", α = 0) mutation profile in ADNI; (B) the mutation profile in ADNI smoothed through a randomised version of the hippocampus network; (C) the mutation profile in ADNI smoothed through a non-brain-related network (umbilical cord). No genes were selected in any of these negative controls with probability higher than 80%.

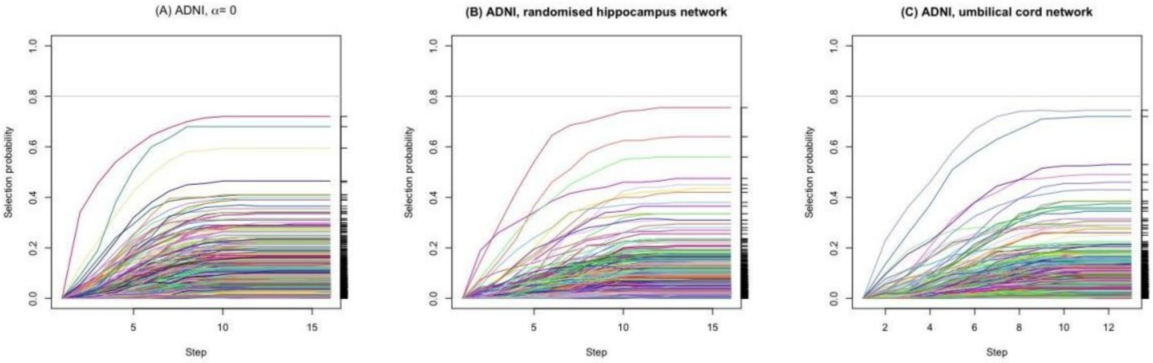

**Figure S4** - Stability selection on a gene burden including only rare deleterious stop-gain, stop-loss and frameshift mutations, smoothed through (A) the hippocampus network from Greene et al (2015) (13,616 genes tested); (B) the STRING PPI network (13,385 genes tested). No genes were selected with probability higher than 80% in either of these alternative scenarios.

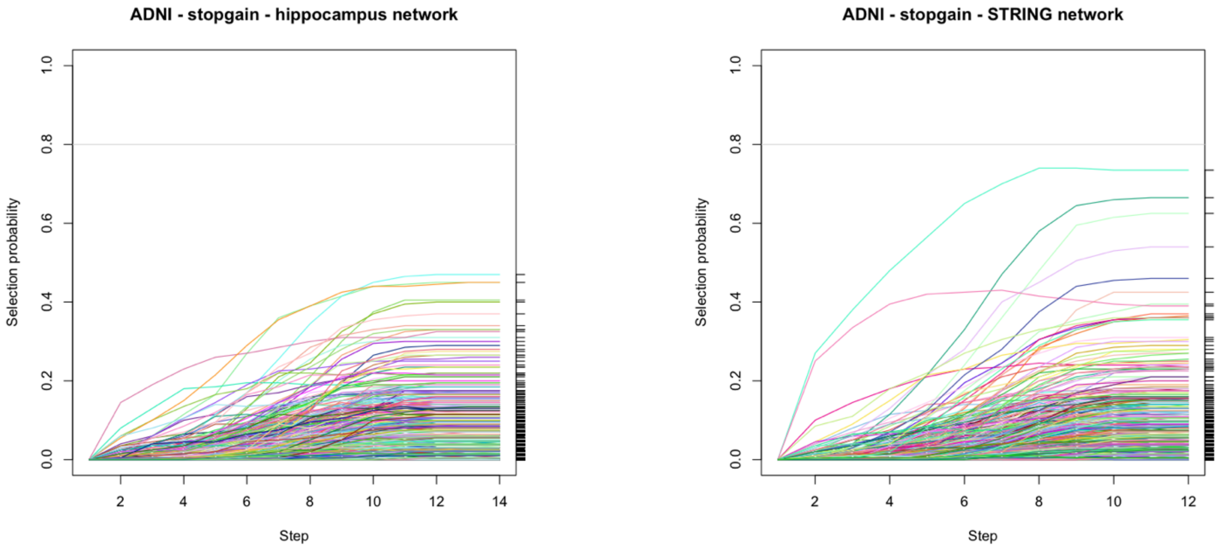

**Figure S5 -** Randomisation control over the amount of overlap between NETPAGE-selected genes in ADSP and the interactome of PFAS. Eighteen out of 29 genes selected in ADSP were observed in the interactome of PFAS (red vertical line). We formed 10,000 replicates of 29 genes selected randomly from the 16,298 genes tested, and counted how many genes in these replicates were also present in the interactome of PFAS. The amount of overlap observed with the genes resulting from the ADSP experiment could not have been achieved by chance, but suggests the presence of functional connections linking NETPAGE's results in the two independent datasets.

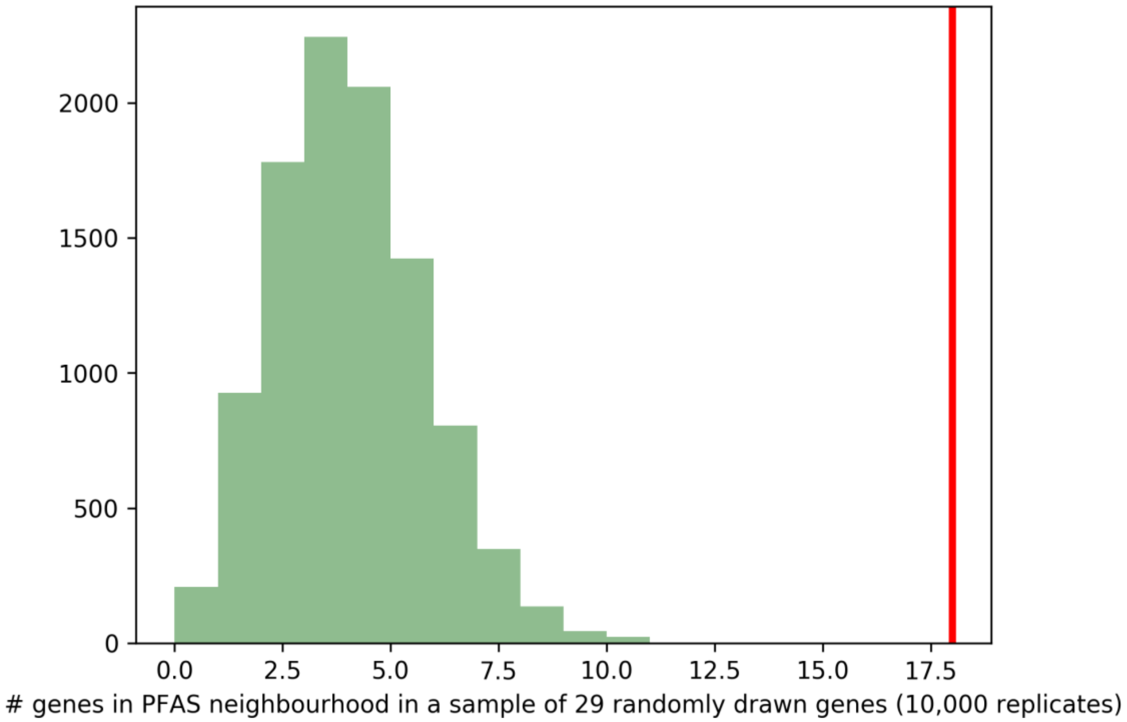

**Figure S5** - Results of gene-based rare variant association testing in ADNI: (A) SKAT-O and (B) mass-univariate testing of smoothed scores against case-control status, in 439 Caucasian participants. Both tests were performed correcting for sex, age, years of education, number of *APOE* 4 alleles, and first two principal components population substructure.

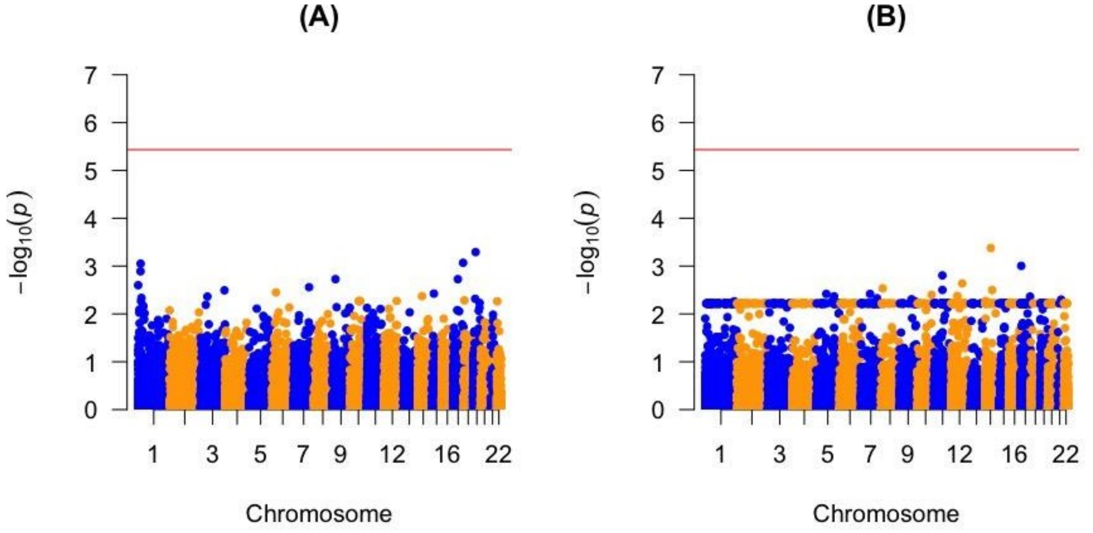

**Figure S6** - Results of gene-based rare variant association testing (SKAT-O) in 10,186 unrelated individuals of Caucasian ancestry from ADSP (16,630 genes tested). The gene-wide significance threshold (red line) was set at 0.05/16,630 = $3x10^{-6}$.

It is interesting to note that SKAT-O on ADSP did not identify any significant gene, in contrast to what reported by [2], despite similar selection criteria for SNV inclusion (particularly the CADD threshold). We are inclined to interpret this result as a consequence of different annotation pipelines and our selection criteria being somewhat stricter, leading to a drastic reduction in the number of SNVs to be grouped and tested (270,165 here vs 918,053 variants in Bis et al. 2018), and likely to result in much lower burden test statistics. Unexpectedly, not even TREM2 showed a significant association in our SKAT-O analysis (p=0.34).

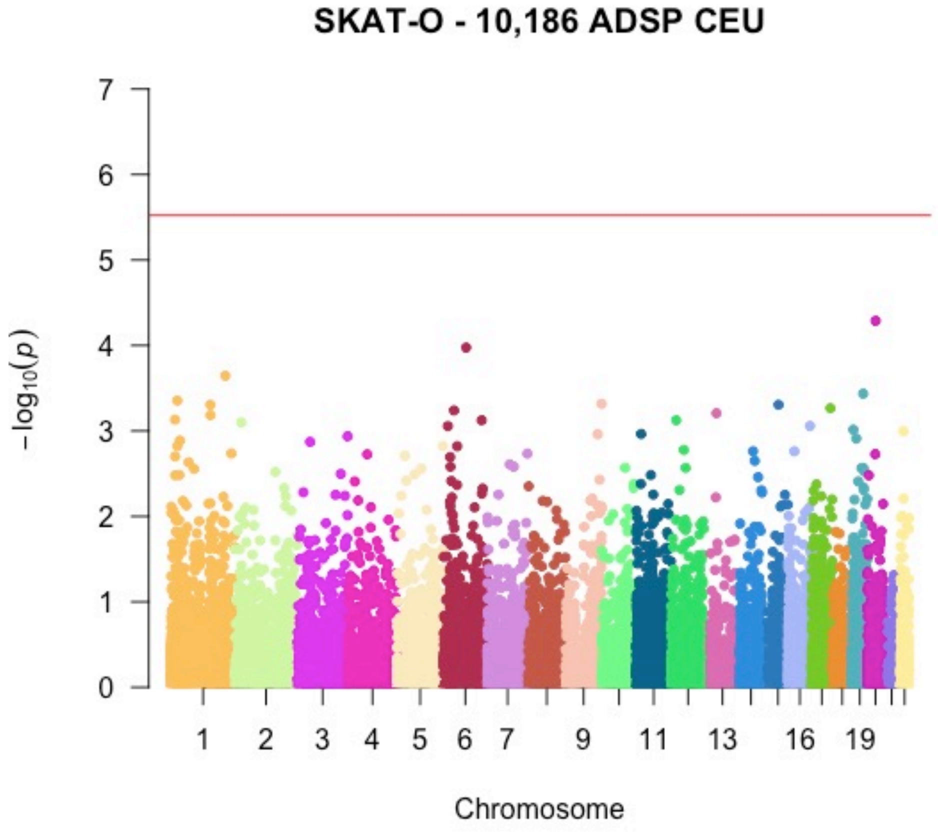

**Figure S7** - Survival analysis for the gene resulting from stability selection in ADNI. (A) Kaplan-Meier survival probability curves stratified by mutation status for *PFAS*. (B) The forest plot depicts hazard ratios from Cox proportional hazards model with confidence intervals and statistical significance. There was still no association between mutation status for PFAS and risk of conversion after covariate correction. (C) Results of the same Cox model fitting as in Figure 4, conducted after removing ADNI subjects diagnosed as cognitively normal at the latest time point available. Therefore the only samples where the conversion event did not occur used in this model were individuals with a stable diagnosis of MCI at the latest time point available. This aimed at partially avoiding circular analysis issues, as cognitively normal individuals were used as control samples in the discovery phase involving stability selection (Figure 3). The score resulting from network propagation for *PFAS* was still seen to be significantly associated with lower risk of conversion to AD after restricting to stable MCI the samples where the conversion event did notoccur.

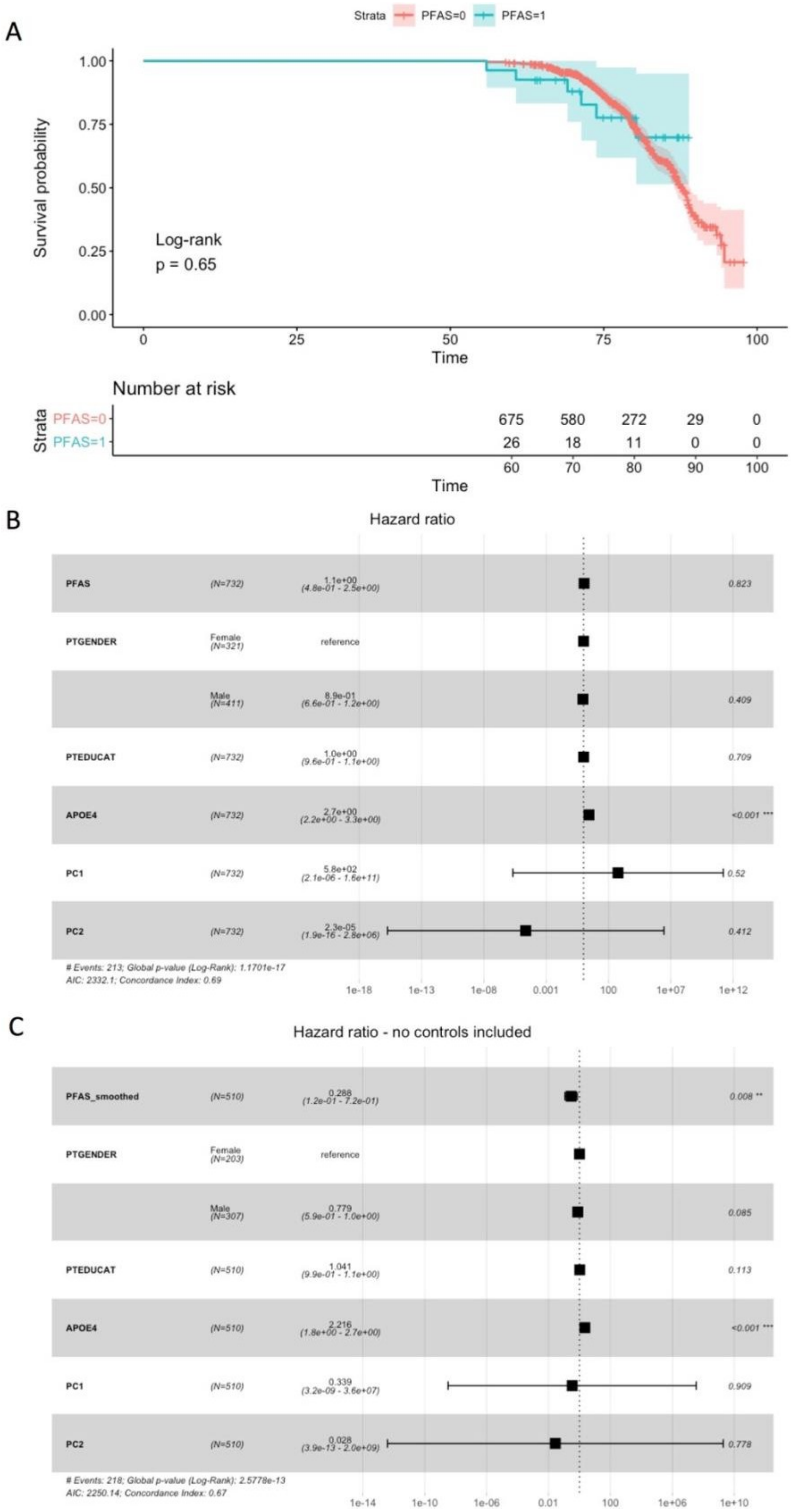
A
Strata
PFAS=0
PFAS=1
Survival probability
1.00
0.75
0.50
0.25
0.00
Log-rank
p = 0.65
0
25
50
75
100
Time
Number at risk
Strata
PFAS=0
PFAS=1
675 580 272 29 0
26 18 11 0 0
60 70 80 90 100
Time
B
Hazard ratio
PFAS (N=732) 1.1e+00 (4.8e-01 - 2.5e+00) 0.823
PTGENDER Female (N=321) reference
Male (N=411) 8.9e-01 (6.6e-01 - 1.2e+00) 0.409
PTEDUCAT (N=732) 1.0e+00 (9.6e-01 - 1.1e+00) 0.709
APOE4 (N=732) 2.7e+00 (2.2e+00 - 3.3e+00) <0.001 ***
PC1 (N=732) 5.8e+02 (2.1e-06 - 1.6e+11) 0.52
PC2 (N=732) 2.3e-05 (1.9e-16 - 2.8e+06) 0.412
### Events: 213; Global p-value (Log-Rank): 1.1701e-17
AIC: 2332.1; Concordance Index: 0.69
1e-18
1e-13
1e-08
0.001
100
1e+07
1e+12
C
Hazard ratio - no controls included
PFAS_smoothed (N=510) 0.288 (1.2e-01 - 7.2e-01) 0.008 **
PTGENDER Female (N=203) reference
Male (N=307) 0.779 (5.9e-01 - 1.0e+00) 0.085
PTEDUCAT (N=510) 1.041 (9.9e-01 - 1.1e+00) 0.113
APOE4 (N=510) 2.216 (1.8e+00 - 2.7e+00) <0.001 ***
PC1 (N=510) 0.339 (3.2e-09 - 3.6e+07) 0.909
PC2 (N=510) 0.028 (3.9e-13 - 2.0e+09) 0.778
### Events: 218; Global p-value (Log-Rank): 2.5778e-13
AIC: 2250.14; Concordance Index: 0.67
1e-14
1e-10
1e-06
0.01
100
1e+06
1e+10

**Figure S8 -** Reduction and visualisation in semantic space, through ReViGO, of the 928 GO terms significantly enriched (at $p_{FDR} < 0.05$) among the 1,449 first and second neighbours of PFAS in the hippocampus network. Top, ontology terms from GO Biological Process; middle, terms from GO Cellular Component; bottom, terms from GO Molecular Function. Bubble color indicates the term p-value (colorbar in lower right-hand corner); size indicates the frequency of the GO term in the underlying database (bubbles of more general terms are larger).

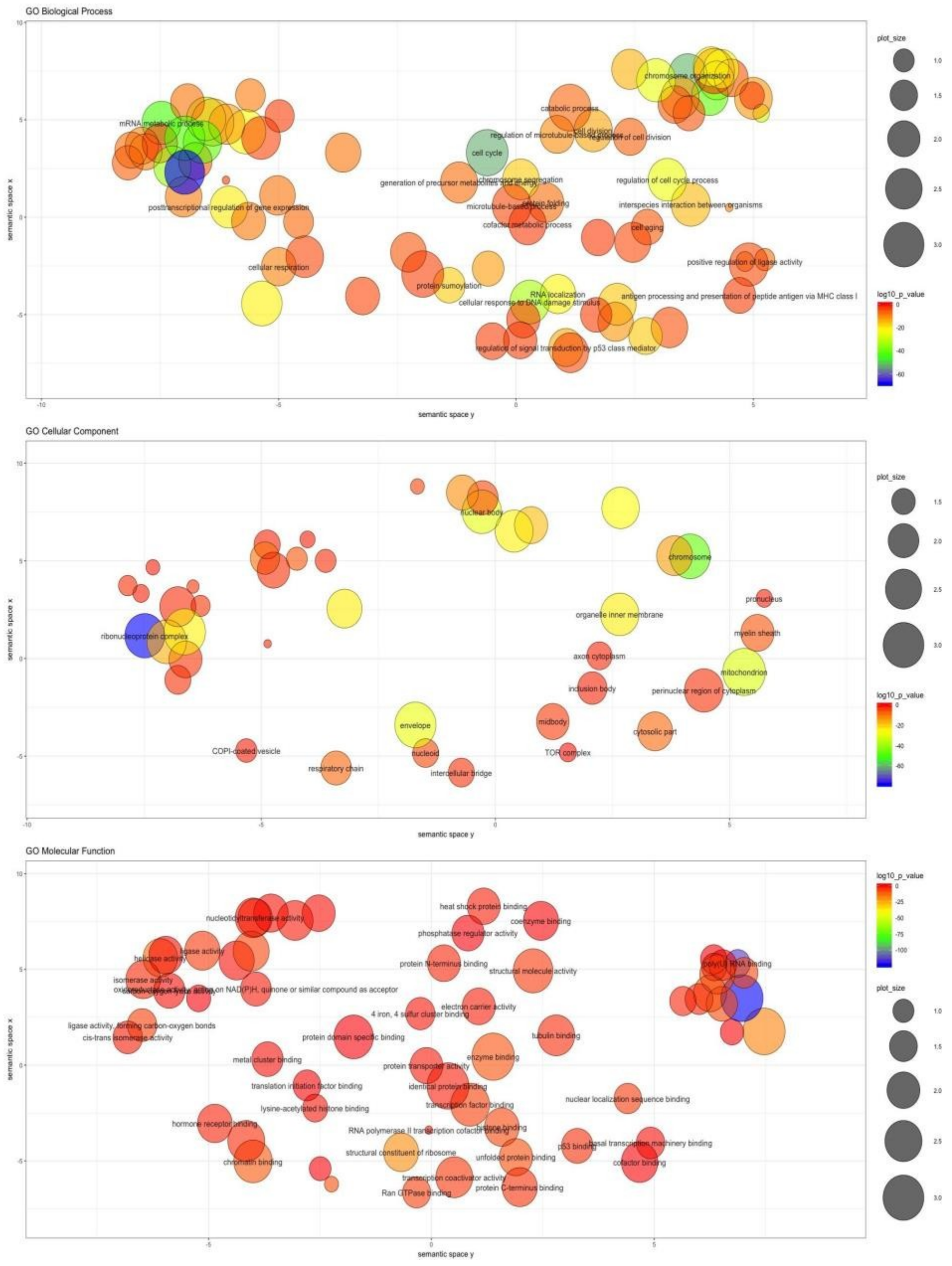

**Figure S9** - Distributions of enrichment p-values for the seven AD-related CGP gene sets, and 1000 gene sets randomly sampled from the background; the red line indicates the location of the non-randomised, uncorrected p-values (P column in Table S2). We show that the significant overlap seen between our genes of interest and the AD-related gene sets curated by Blalock (Table S2) could not be achieved by chance, as none of the 1000 randomly drawn gene sets achieved smaller p-values.

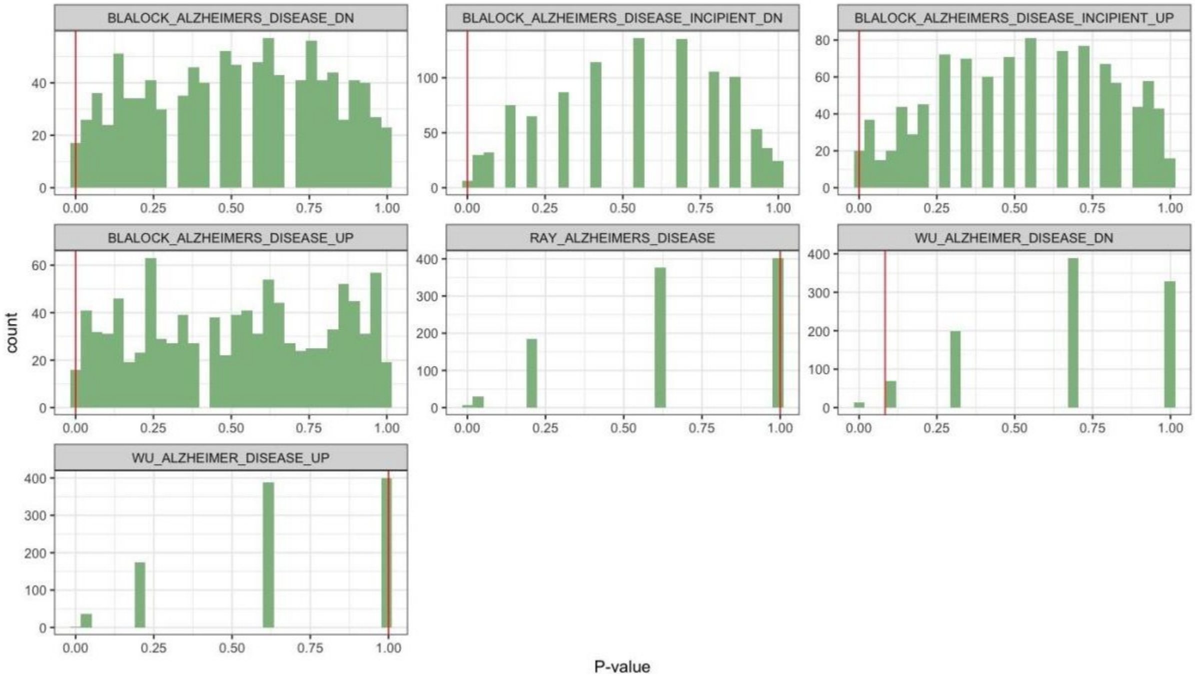

**Figure S10 -** Differential expression analysis for 30 selected genes in the Mount Sinai Brain Bank parahippocampal gyrus expression dataset. Full numerical results for all pairwise comparisons and their significance are provided in Supplementary Table 3.

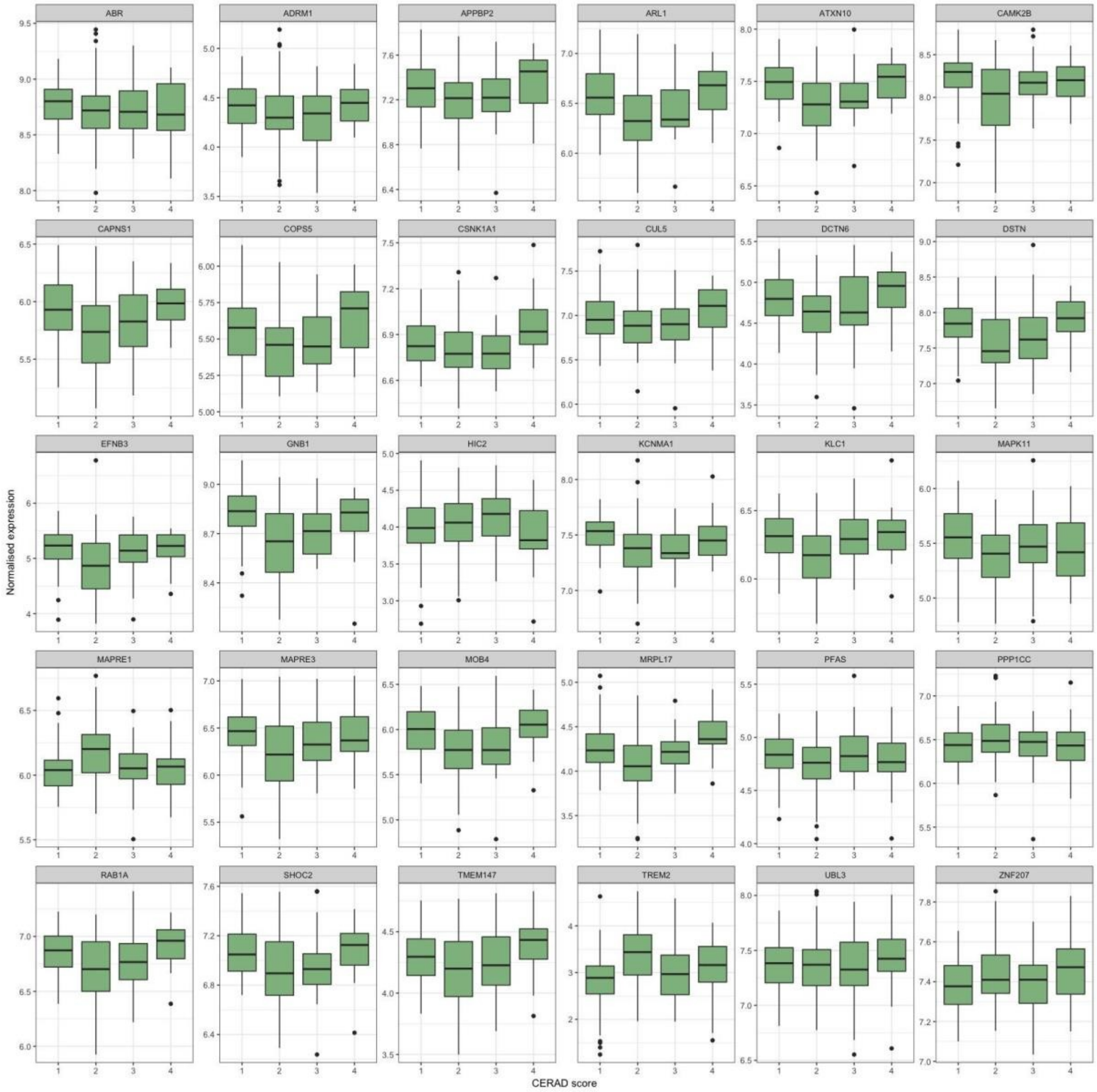

**Figure S11 -** Randomisation control over the differential expression analysis in the Mount Sinai Brain Bank dataset. We formed 1,000 sets of 30 randomly sampled genes and tested each of them for pairwise differences in expression levels among CERAD categories. We then counted how many genes in each set showed significantly different expression in at least one comparison and plotted their distribution. The red line indicates the 16 genes that showed significantly different expression in the original gene set of interest (i.e., the 30 genes selected in ADNI and ADSP). The data clearly shows that this amount of dysregulation could not be observed by chance, but is indeed linked to the overrepresentation of disease-related genes in the set of interest.

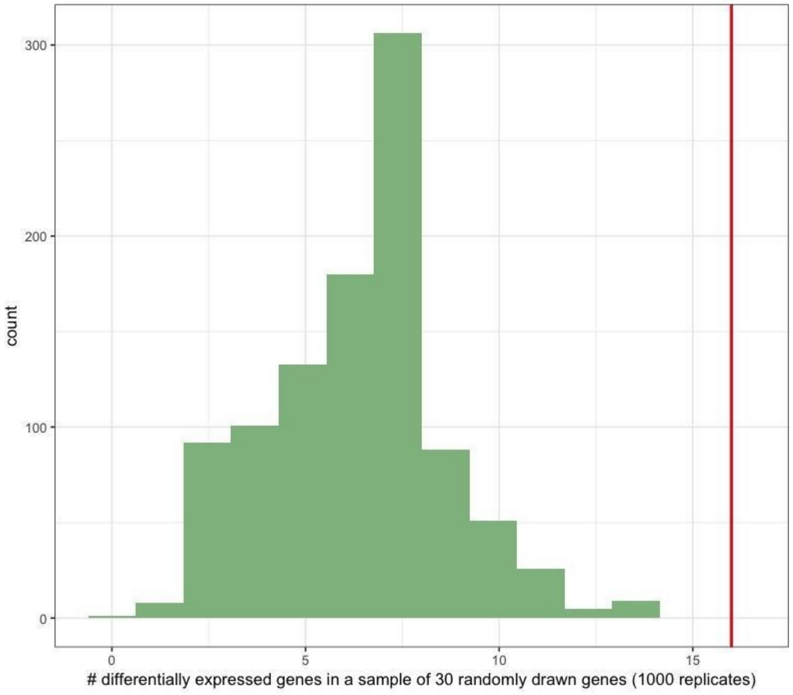

#### References


1. Saykin AJ, et al. (2015) Genetic studies of quantitative MCI and AD phenotypes in ADNI: Progress, opportunities, and plans. *Alzheimer's Dement* 11(7). doi:10.1016/j.jalz.2015.05.009.
2. Li H, Durbin R (2009) Fast and accurate short read alignment with Burrows-Wheeler transform. *Bioinformatics*. doi:10.1093/bioinformatics/btp324.
3. Depristo MA, et al. (2011) A framework for variation discovery and genotyping using next-generation DNA sequencing data. *Nat Genet*. doi:10.1038/ng.806.
4. Chang CC, et al. (2015) Second-generation PLINK: rising to the challenge of larger and richer datasets. *Gigascience* 4(1):7.
5. Greene CS, et al. (2015) Understanding multicellular function and disease with human tissue-specific networks. *Nat Genet* 47(6):569–76.
6. Hagberg AA, Schult DA, Swart PJ (2008) Exploring network structure, dynamics, and function using NetworkX. *Proceedings of the 7th Python in Science Conference (SciPy)* doi:10.1016/j.jelectrocard.2010.09.003.
7. Hofree M, Shen JP, Carter H, Gross A, Ideker T (2013) Network-based stratification of tumor mutations. *Nat Methods* 10(11):1108–1115.
8. Allen M, et al. (2016) Human whole genome genotype and transcriptome data for Alzheimer's and other neurodegenerative diseases. *Sci Data* 3:160089.
9. Wang M, et al. (2018) The Mount Sinai cohort of large-scale genomic, transcriptomic and proteomic data in Alzheimer's disease. *Sci Data* 5:180185.
10. Mirra SS, et al. (1991) The Consortium to Establish a Registry for Alzheimer's Disease (CERAD). Part II. Standardization of the neuropathologic assessment of Alzheimer's disease. *Neurology* 41(4):479–86.
